# SFTSV NSs protein is a novel tick antiviral RNAi response suppressor

**DOI:** 10.1101/2025.08.14.670292

**Authors:** Mazigh Fares, Melanie McFarlane, Rhys H. Parry, Rozeena Arif, Andrew T. Clarke, Wael Kamel, Kelsey Davies, Riona Datta-Savage, Ulrich Schwarz-Linek, Lesley Bell-Sakyi, Marine J. Petit, Esther Schnettler, Alfredo Castello, Alain Kohl, Benjamin Brennan

**Author notes:** joint first authors (Mazigh Fares, Melanie McFarlane, Rhys H. Parry).

## Abstract

Severe fever with thrombocytopenia syndrome virus (SFTSV) is an emerging tick-borne phenuivirus causing high mortality in humans. While the non-structural protein NSs is dispensable for replication in interferon-deficient mammalian cells, we demonstrate for the first time that NSs is essential for viral replication in tick cells. SFTSV infection triggers canonical Dicer-2-mediated antiviral RNA interference (RNAi) in tick cells, producing virus-derived small interfering RNAs (siRNAs) that target viral transcripts for degradation. We show that NSs functions as a viral suppressor of RNAi by selectively engaging and depleting single-stranded RNAs derived from 22-nucleotide siRNAs, likely limiting their incorporation into RNA-induced silencing complexes (RISC). Complementation with a heterologous RNAi suppressor (p19 protein) partially rescues replication of NSs-deficient virus, validating the RNAi suppressive function of NSs. These findings reveal that successful tick-borne viral replication requires host-specific immune evasion strategies and establish NSs-mediated RNAi suppression as essential for SFTSV persistence in arthropod vectors.

**Significance Statement:** This study reveals a critical requirement for SFTSV infection that differs between hosts. While the viral NSs protein is dispensable in interferon deficient mammalian cells, it is essential for replication in tick cells, the natural vectors of the virus. We show that NSs functions as a viral RNA silencing suppressor by associating with virus derived small RNAs and limiting their availability to the tick antiviral RNA interference machinery. This provides mechanistic insight into how tick-borne viruses evade arthropod immune defences through engagement of functional siRNAs rather than broad inhibition of RNA interference pathways. Together, these findings demonstrate that SFTSV employs distinct immune evasion strategies in different hosts and identify NSs as a key determinant of vector cell infection.

## MAIN TEXT

Severe fever with thrombocytopenia syndrome virus (SFTSV; *Bandavirus dabieense*; *Phenuiviridae*) emerged as a significant threat to human health following its discovery in central China in 2009 (1), where it was identified as the causative agent of a severe haemorrhagic fever syndrome. This tick-borne phenuivirus rapidly gained attention due to its high case fatality rate of 6-30%. Its distinctive clinical presentation is characterized by fever, thrombocytopenia, leukocytopenia, and gastrointestinal manifestations (2). The geographic range of SFTSV has since expanded across East Asia, with confirmed cases reported in South Korea, Japan, Taiwan, and Vietnam, establishing it as a regional public health concern of growing importance (3). Currently, no effective antiviral drugs or vaccines exist to treat SFTS disease.

SFTSV transmission occurs primarily through the bite of infected ixodid ticks, with several species identified as potential vectors (4). *Haemaphysalis longicornis* serves as the principal vector across much of the SFTSV geographical range, while *Haemaphysalis flava* and *Rhipicephalus microplus* may also contribute to transmission cycles in specific regions (5). Ticks acquire SFTSV through blood feeding on infected vertebrate hosts and can subsequently transmit the virus to humans and other susceptible animals. This complex ecology involves multiple tick species and some amplifying vertebrate hosts (hedgehogs and small rodents), creating diverse transmission pathways that complicate control efforts (6).

Arthropod vectors, including ticks, have evolved sophisticated innate immune mechanisms to combat viral infections, with RNA interference (RNAi) representing a primary antiviral defence pathway (7, 8). Unlike mammals, in which type I interferon responses dominate antiviral immunity (9), arthropods rely heavily on RNAi machinery to recognise and degrade viral nucleic acids. Tick antiviral RNAi responses produce characteristic 22-nucleotide small interfering RNAs (siRNA) through Dicer-2 processing of viral double-stranded RNA intermediates, which are subsequently incorporated into RNA-induced silencing complexes (RISC) to target complementary viral transcripts for degradation (8, 10). Distinguishing authentic virus-derived siRNAs (vsiRNA) from RNA degradation products requires methylation-sensitive approaches, as functional siRNAs undergo Hua Enhancer-1 (HEN1)-mediated 2’-O-methylation upon RISC loading (11–15). The effectiveness of arthropod RNA silencing as an antiviral mechanism has been demonstrated across diverse vector species (flies and mosquitoes) (16), yet our understanding of how tick-borne viruses evade or suppress these responses remains limited (17).

The evolutionary arms race between viruses and RNAi defenses has driven many viruses to evolve suppressors of RNAi (VSRs) that counteract antiviral silencing (18, 19). They employ diverse mechanisms to inhibit RNAi activity, including sequestration of siRNAs, interference with Dicer processing, and disruption of RISC assembly and function. While NSs proteins from plant-infecting tospoviruses are well-characterized RNAi suppressors that bind double-stranded siRNAs (20), vertebrate-infecting bunyaviruses show variable suppressor activities (21). Crucially, no arthropod-specific RNAi suppression mechanisms have been definitively established for tick-borne phenuiviruses.

Despite SFTSV’s emergence as a significant public health threat and evidence of vsiRNA production in infected ticks (22), the specific mechanisms by which tick-borne phenuiviruses evade or suppress arthropod RNAi responses remain unknown. While the non-structural protein (NSs) of SFTSV has been suggested as a possible RNAi suppressor (22, 23), its actual role and mechanism of action in arthropod cells have yet to be definitively established.

Here, we demonstrate for the first time that SFTSV exhibits fundamentally different replication requirements in tick versus mammalian cell cultures, with NSs proving essential for viral maintenance in tick cells. Using comprehensive molecular analyses including methylation-sensitive small RNA sequencing, structural modelling, and complementation studies with heterologous suppressors, we reveal that NSs functions as a unique viral suppressor of RNAi by selectively sequestering the pool of virus-derived single stranded siRNAs away from the RNAi machinery, thereby enabling productive infection in tick cells and enhancing vector-borne transmission potential.

## RESULTS

### SFTSV infects cell lines derived from tick species reported to act as competent vectors

To identify suitable tick cellular models for SFTSV infection, we tested multiple cell lines representing both *Metastriata* (*Rhipicephalus* and *Hyalomma* spp.) and *Prostriata* (*Ixodes* and *Amblyomma* spp.) lineages of the family *Ixodidae*, using wildtype (wt) SFTSV strain HB29 (SI Appendix, Fig. S1) (1, 24). Viral yield assays revealed SFTSV efficiently replicated in cell lines derived from *Rhipicephalus* and *Hyalomma* spp, evidenced by increased viral titres and nucleocapsid protein detection in infected lysates (SI Appendix, Fig. S1A,B). SFTSV replication was minimal in phylogenetically distinct *Ixodes* and *Amblyomma* spp. tick cell lines, despite these species vectoring other members of the *Phenuiviridae* such as Uukuniemi (25) and Heartland viruses (26, 27). Given that bunyaviruses establish non-cytolytic, continuous viral production in tick cells (25, 28), we monitored infectious virus production daily over 10 days at low MOI (SI Appendix, Fig. S1C). Viral titres steadily decreased over time in *Ixodes* spp. cell lines, but continuous production was evident in *Rhipicephalus* spp. cultures. After 10 days post-infection (dpi), all cultures were split 1:2 and monitored for 14 days; clear SFTSV persistence in BME/CTVM6 and BME/CTVM23 lines (SI Appendix, Fig. S1D) confirms their utility as robust models for studying SFTSV arthropod cell infection (29). These results suggest SFTSV entry and replication may depend on host cellular factors that vary among tick species and demonstrate that permissive cell lines can maintain viable virus over extended culture periods.

### SFTSV NSs protein expression is repressed in tick cells yet is essential for sustained virus replication

We previously established that SFTSV NSs is a potent interferon antagonist (30) and virulence factor in animal models (31–33). While NSs is essential for replication in interferon-competent mammalian cells, it is dispensable in IFN-deficient cell lines (34), suggesting cell-type-specific functions beyond IFN antagonism. However, the molecular basis for NSs function in tick cells, the natural arthropod vector, has remained unexplored. To address this knowledge gap, we investigated NSs function during SFTSV infection of tick cells. We infected BME/CTVM6 cells, selected for their robust and sustained wt SFTSV replication (SI Appendix, Fig. S1), with either wt or NSs-deleted (delNSs) SFTSV across a range of multiplicities of infection. We observed that wt SFTSV demonstrated stable and abundant production of infectious virus throughout the experiment, with SFTSV titres increasing by approximately 3-log_10_ FFU/ml over the time course (Fig. 1A). In contrast, delNSs SFTSV failed to maintain infectious progeny, with viral titres declining progressively over the infection time course (Fig. 1B). Consistent with this, Western blot analysis of delNSs SFTSV-infected cell lysates revealed no detectable nucleocapsid protein, indicating complete failure to establish a productive infection in tick cells (Fig. 1C). Notably, NSs protein was undetectable in cells infected with both wt and delNSs SFTSV, despite this protein being detected in mammalian cell controls (Vero E6). This observation was unexpected given the apparent requirement for NSs in tick cells. To investigate this paradox further, we performed immunofluorescence microscopy on infected BME/CTVM6 cells (Fig. 1D). Quantitative analysis revealed that delNSs SFTSV infection led to only 2-8% nucleocapsid-positive cells (Fig. 1E,F). In contrast, 60-80% of cells were positive for nucleocapsid protein following wt SFTSV infection across all tested MOIs, demonstrating efficient viral replication (Fig. 1G,H).

**Fig. 1.**
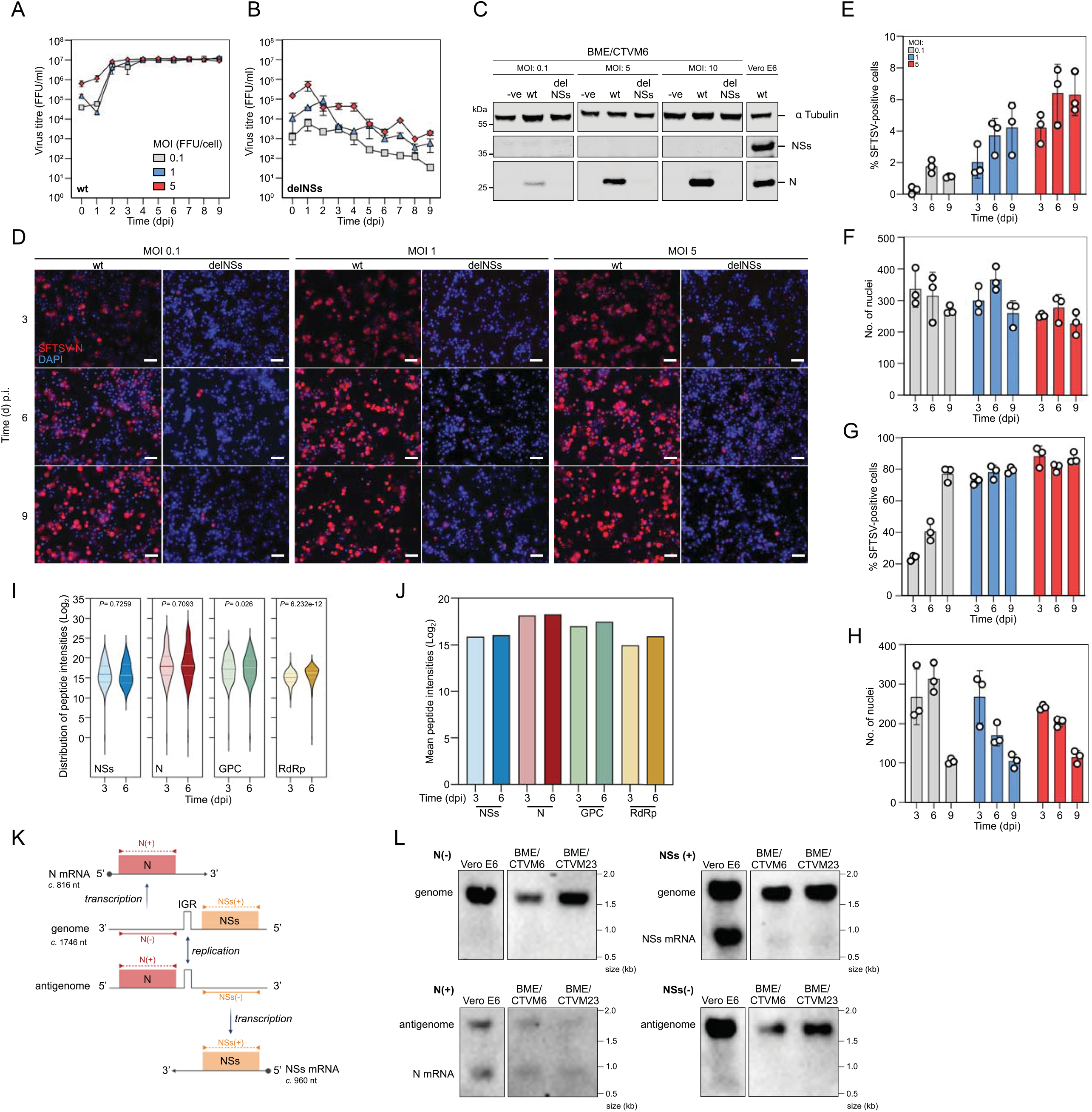
Initial characterisation of wt SFTSV replication in a diverse range of tick cells. **(A-B)** Titre of **(A)** wt or **(B)** delNSs SFTSV in the supernatant of infected BME/CTVM6 cells (MOI: 0.1 [grey]; 1 [blue]; or 5 [red]) assayed at the indicated timepoints p.i. Data are plotted as the mean virus titre (FFU/ml) ± SD of n=2 biological replicates. **(C)** Representative images of cell lysates derived from cell monolayers in **(A)** and **(B)** harvested at 72 h p.i. As a control, Vero E6 cells were infected with wt SFTSV (MOI: 1) and cell lysates were prepared at 72 h p.i. Western blots were probed with anti-tubulin (loading control), anti-SFTSV NSs or anti-SFTSV N antibodies to detect cellular and viral protein expression. ‘-ve’: mock-infected cell lysate, ‘wt’: wt SFTSV infected cell lysate, ‘delNSs’: delNSs SFTSV infected cell lysate. Data show a representative experiment (n=3). **(D)** BME/CTVM6 cells seeded onto coverslips were infected with wt- or delNSs-SFTSV at the indicated MOI and were imaged at the timepoints indicated using DAPI (blue) and anti-SFTSV N (red). Imaging was carried out using a Zeiss LSM 880 Confocal Microscope, scale bar is equal to 50 μm. **(E)** Percentage of BME/CTVM6 cells infected with delNSs SFTSV (MOI: 0.1 [grey]; 1 [blue]; or 5 [red]) at the timepoints indicated, evidenced by positive staining with anti-SFTSV N antibody. Data are plotted as the mean percent SFTSV-positive cells ± SD of n=3 randomised fields of view. **(F)** Number of nuclei enumerated per field of view in (E). Data are plotted as the mean number of nuclei observed ± SD of n=3 randomised fields of view. **(G)** Percentage of BME/CTVM6 cells infected with wt SFTSV (MOI: 0.1 [grey]; 1 [blue]; or 5 [red]) at the timepoints indicated, evidenced by positive staining with anti-SFTSV N antibody. Data are plotted as the mean percent SFTSV-positive cells ± SD of n=3 randomised fields of view. **(H)** Number of nuclei enumerated per field of view in (G). Data are plotted as the mean number of nuclei observed ± SD of n=3 randomised fields of view. **(I)** Violin plot showing peptide intensity distributions (y-axis: Log_2_) for wt SFTSV-infected BME/CTVM6 cells at 3- and 6-dpi. Examining peptides derived from non-structural protein (NSs; AJD86041), nucleocapsid protein (N; AJD86040), glycoprotein precursor (GPC; AJD86039) or RNA-dependent RNA polymerase (RdRp; AJD86038). Data taken from n=4 biologically independent replicates. **(J)** Bar Chart showing mean peptide intensities (y-axis: Log_2_) for same data set presented in (I). **(K)** Schematic of SFTSV S genomic RNA configuration showing DIG-labelled probe binding regions used in (L). Probes detect viral genomic (−) or antigenomic (+) sense RNA as indicated. Double-ended arrows indicate genomic replication; single-ended arrows indicate transcription. Black circles indicate 5′ cap structures on mRNAs. **(L)** Northern blot analysis of viral RNAs in infected cells. BME/CTVM6 and BME/CTVM23 cells were infected with wt SFTSV (MOI: 0.1) and total RNA was extracted at 6 dpi. Vero E6 cells infected with wt SFTSV were used as a positive control (RNA extracted at 72 h p.i.). DIG-labelled probes complementary to N or NSs coding sequences detected viral genomic (−) or antigenomic (+) sense RNAs. RNA sizes are indicated on the right.

To gain a deeper understanding of the viral protein abundance in SFTSV infected tick cells we performed proteomic analysis of BME/CTVM6 cells infected with wild-type SFTSV at 3- and 6-dpi, examining both the distribution of peptide intensities (Fig. 1I) and mean peptide intensities (Fig. 1J) for viral proteins NSs, nucleocapsid protein (N), glycoprotein precursor (GPC), and RNA-dependent RNA polymerase (RdRp). The peptide intensity distributions for NSs remained unchanged between time points, while N showed a non-significant increase (*P =* 0.709). This reflects that N protein peptides already exhibited high intensities and broad distribution at 3 dpi, suggesting a detection plateau had been reached. In contrast, GPC and RdRp exhibited significant increases (*P =* 0.026 and *P =* 6.23 × 10^-12^, respectively), indicative of enhanced viral replication and assembly processes (Fig. 1I). Analysis of mean peptide intensities revealed temporal increases for N, GPC, and RdRp from 3 to 6 dpi, while NSs maintained the lowest expression levels with minimal temporal change (Fig. 1J). Notably, while NSs-derived peptides were detectable by mass spectrometry, NSs protein intensity levels remained below the detection threshold of Western blot analysis (Fig. 1C), suggesting tightly regulated, low-level expression of this protein. This differential sensitivity between proteomic and immunoblot detection methods reveals that NSs is present but maintained at substantially lower levels compared to structural and replication proteins during SFTSV infection of tick cells.

### NSs subgenomic mRNA expression is reduced in SFTSV-infected tick cells compared to mammalian cells

The SFTSV NSs protein is encoded within the viral S segment, which employs an ambisense coding strategy to produce both NSs and the N protein, consistent with all phenuiviruses (35). The NSs open reading frame is translated from a subgenomic mRNA transcribed from the antigenomic S RNA template following viral genome replication. To investigate the molecular basis for NSs repression in tick cells, we analysed viral RNA species derived from the S segment during infection of two *R. microplus* tick cell lines and compared these results to mammalian Vero E6 cells as controls.

We employed Northern blot analysis using strand-specific RNA probes to target either the N or NSs coding regions and to distinguish genomic from antigenomic RNA species (Fig. 1K). Productive viral replication was confirmed in wild-type SFTSV-infected tick cells through detection of both genomic [detected by N(−) and NSs(+) probes] and antigenomic [detected by N(+) and NSs(−) probes] S segment RNA at 6 dpi. While N mRNA was weakly but reproducibly detectable in both tick cell lines and mammalian cell controls, NSs subgenomic mRNA levels were markedly reduced in tick cells compared with Vero E6 cells when probed with the NSs(+) probe (Fig. 1L).

The observed reduction in NSs subgenomic mRNA levels in tick cells provides a plausible explanation for the diminished NSs protein expression during infection. However, the underlying mechanism, whether due to transcriptional attenuation or altered RNA stability, remains unresolved. While our Northern blot data confirm reduced mRNA accumulation, further investigation is needed to determine whether this reflects regulated transcription termination, selective degradation or post-transcriptional silencing. This finding also raises important questions about how NSs functions during early infection, before its apparent repression.

### SFTSV infection triggers small RNA production in tick cells

Given that tick cells rely on RNAi as a primary antiviral defence mechanism, we next examined whether wt SFTSV replication, and specifically NSs mRNA, is targeted by the tick RNAi pathway. To determine whether the observed reduction in SFTSV NSs mRNA reflected targeted RNAi-mediated degradation, we used beta elimination to distinguish between RISC-loaded siRNAs, unloaded siRNAs, and RNA degradation products. Small RNA samples were therefore subjected to beta elimination prior to sequencing analysis. This treatment selectively removes degradation fragments and non-methylated siRNAs containing 2’,3’-cyclic phosphate termini while preserving HEN1-methylated, RISC-loaded siRNAs bearing 5’-phosphate and 3’-hydroxyl groups (11–15). Since both degradation products and non-RISC-loaded siRNAs exhibit similar size distributions (20-24 nt), protection by HEN1 methylation during beta elimination enables unambiguous identification of functional vsiRNAs and confirms RNAi pathway activation against SFTSV (12, 13).

The effectiveness of beta elimination was validated in uninfected BME/CTVM6 cells (SI Appendix, Fig. S2A), the Ago2-deficient *Aedes aegypti* AF525 cell line (SI Appendix, Fig. S2B), or Semliki Forest virus (SFV)-infected AF525 cells (SI Appendix, Fig. S3A,B) by published methodologies, noting that siRNAs produced by mosquito cells are 21nt in length (11). Following SFTSV infection of BME/CTVM6 cells, approximately 10^5^ reads within the small RNA size range (18-30 nt) mapped to each SFTSV genomic segment in untreated samples, with read abundance increasing between 3- and 6-dpi. Although beta elimination reduced overall read counts, the proportional increase in reads mapping to SFTSV RNAs from 3 to 6 dpi was preserved, indicating sustained small RNA responses targeting SFTSV throughout infection (Fig. 2A).

**Fig. 2.**
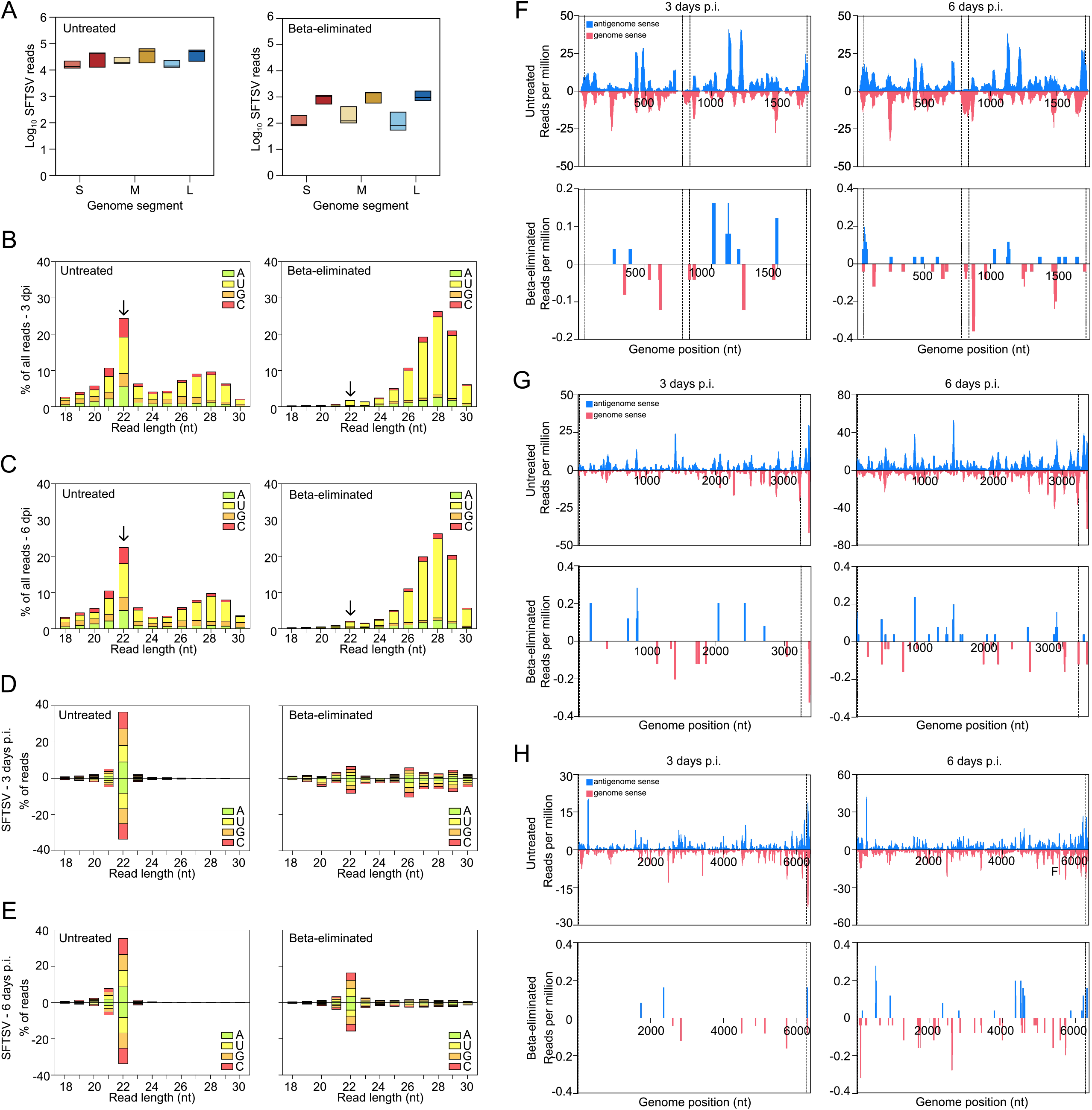
SFTSV infection of tick cells triggers canonical antiviral RNA interference (RNAi) and production of 22-nt virus-derived small interfering RNAs (vsiRNAs). BME/CTVM6 cells were infected with wt SFTSV (MOI: 0.1) and total cellular RNA was extracted from infected cells at 3 (light shade) or 6 (dark shade) dpi. Following RNA extraction, samples were either left untreated or processed in a beta-elimination assay, then subjected to small RNA sequencing. β-elimination selectively removes non-methylated RNA fragments while leaving 2’-O-methylated, RISC-loaded vsiRNAs intact, enabling functional siRNAs to be distinguished from degradation products. **(A)** Total number of small RNA reads (18-30 nt) in infected cells at 3 (light shading) and 6 (dark shading) dpi. mapping to each wt SFTSV genome segment in untreated samples (left) or beta-eliminated samples (right). SFTSV genome segments: S (KP202165), M (KP202164), L (KP202163). An increase in viral small RNA reads indicates activation of antiviral RNAi over time. **(B)** Histogram of all small RNA reads (18-30nt) from untreated (left) or beta-eliminated (right) samples from infected cells at 3 dpi. Read counts are coloured by the identity of the 5′ nucleotide and grouped by read length. Data are shown as the mean percentage of total mapped reads (y-axis); representative of n=3 biologically independent samples. **(C)** As per (B), but RNAs were extracted from wt SFTSV-infected cells at 6 dpi. At 6 dpi, a pronounced 22-nt peak re-emerges, indicating accumulation of methylated vsiRNAs that have entered Argonaute-containing RISC complexes. **(D)** Histogram of small RNA reads (18-30 nt) from untreated (left) or beta-eliminated (right) samples, mapped to the combined SFTSV antigenomic (positive values) and genomic (negative values) strand RNAs in infected cells at 3 dpi. Read counts are coloured by the identity of the 5′ nucleotide and grouped by read length. Data are shown as the mean percentage of total mapped reads (y-axis); representative of n=3 biologically independent samples. **(E)** As per (D), but RNAs were extracted from wt SFTSV-infected cells at 6 dpi. Loss of the 22-nt signal after β-elimination confirms that early vsiRNAs are not yet methylated or RISC-loaded, while its recovery at 6 dpi demonstrates canonical RNAi activation. **(F-H)** Mapping of SFTSV-derived 22 nt vsiRNAs (reads per million) across the **(F)** S, **(G)** M, and **(H)** L genome segments at 3 (left) and 6 (right) dpi in BME/CTVM6 cells. Data show antigenomic (blue) and genomic (pink) sense strands for untreated (top) or beta-eliminated samples (bottom) to remove 2’-O-methylated small RNAs. Numbers indicate nucleotide positions (nt). SFTSV genome segments: S (KP202165), M (KP202164), L (KP202163). Dashed lines represent the boundaries of viral open reading frames and untranslated/intergenic regions. Concentrated vsiRNA peaks within the S segment intergenic region and NSs open reading frame correspond to targeted degradation of viral RNAs by the tick RNAi machinery. Together, these data demonstrate that SFTSV infection activates Dicer-2–mediated processing of viral RNA into 22-nt vsiRNAs, which progressively acquire methylation and RISC loading as infection proceeds.

The 5’ nucleotide composition of all 22 nt vsiRNAs showed strong bias toward uridine and adenine residues, consistent with arthropod Dicer-2 processing specificity (36). Beta elimination treatment virtually eliminated the 22 nt peak at both time points, indicating these vsiRNAs lacked 3’-terminal methylation and remained susceptible to chemical modification (Fig. 2B,C). Analysis of vsiRNAs mapping to SFTSV genome and antigenome sequences at both 3- and 6-dpi revealed pronounced enrichment for 22 nt species in untreated samples, with comparable distributions observed for genomic and antigenomic reads (Fig. 2D,E). Conversely, longer small RNAs (24-30 nt) were largely unaffected by beta elimination, consistent with 2’-O-methylated piRNAs (PIWI-interacting RNAs; these are small non-coding RNAs, typically 24-31 nucleotides long, that bind to PIWI proteins to regulate gene expression) or degradation fragments that escaped chemical treatment.

Notably, the selective loss of Dicer-2-processed 22 nt reads following beta elimination at 3 dpi demonstrates that the majority of SFTSV-derived siRNAs in BME/CTVM6 cells lack methylation modifications, and remain susceptible to chemical modification (Fig. 2D). However, this transient suppression of vsiRNAs from Ago2 is eventually overwhelmed by 6 dpi, evidenced by the strong 22 nt peak of methylated-vsiRNAs in the beta-eliminated sample, indicating a resumption in the targeting of viral RNA sequences by the RNAi machinery (Fig. 2E).

While HEN1-mediated methylation of RISC-loaded siRNAs is conserved from plants to arthropods (12, 37), the dynamics of this process during viral infection in tick cells have not been previously characterized. Our detection of methylated small RNAs in mock-infected controls (SI Appendix, Fig. S2) and their recovery at 6 dpi confirms that BME/CTVM6 cells possess functional HEN1 methylation machinery. Therefore, markedly reduced levels of methylated SFTSV-specific siRNAs at 3 dpi (Fig. 2D), followed by their subsequent accumulation at 6 dpi (Fig. 2E), indicates active viral suppression of RNAi responses during early infection that is eventually overcome by sustained antiviral pressure, enabling initial viral replication. The observed NSs transcript degradation supports this interpretation (Fig. 1L), demonstrating that functional RNAi activity ultimately prevails despite initial viral evasion mechanisms.

### RNAi-mediated vsiRNA targeting exhibits temporal dynamics and segment-specific preferences across the SFTSV genome

Given that 22-nt species constituted the predominant small RNA population (which corresponds to the typical length of vsiRNAs produced in tick cells), we mapped vsiRNA coverage across genomic and antigenomic strands of the SFTSV small (S; Fig. 2F), medium (M; Fig. 2G), and large (L; Fig. 2H) segments at 3- and 6-dpi in untreated and beta-eliminated samples. SFV served as a control for beta-elimination efficacy and mapping specificity (SI Appendix, Fig. S3C,D).

SFTSV vsiRNA accumulation increased substantially between 3- and 6-dpi across all segments, indicating progressive amplification of the RNAi response. In untreated samples, the M and L segments exhibited extensive targeting, with pronounced accumulation at untranslated regions and segment termini. Beta-elimination confirmed sustained SFTSV targeting by the tick RNAi machinery, demonstrating that observed signals represent authentic vsiRNAs rather than 2’-O-methylated small RNA contaminants. Notably, the S segment displayed concentrated targeting within the intergenic region and the NSs open reading frame on the genomic strand, particularly around the areas previously identified as the transcription termination sites for the N and NSs mRNAs (34), indicating RNAi-mediated cleavage of both genomic RNA and NSs subgenomic transcripts. This finding provides a mechanistic explanation for the reduced NSs mRNA detection observed by Northern blot analysis (Fig. 1L). The strand-specific and segment-preferential targeting patterns demonstrate that tick cells mount a sophisticated, temporally regulated antiviral RNAi response against SFTSV infection, which the wt virus can overcome, whereas a NSs-deletant virus cannot.

### SFTSV infection triggers canonical Dicer-2-mediated vsiRNA production in tick cells

To characterise the dsRNA cleavage patterns during SFTSV infection, we analysed small RNA overlap profiles and strand complementarity patterns mapping to viral segments at 3- and 6-dpi (Fig. 3). At both timepoints, viral small RNAs predominantly exhibited 20-22 nucleotide lengths with a clear preference for 20 nucleotide overlaps between complementary strands (Fig. 3A,C). Strand overlap analysis revealed enhanced complementarity at positions resulting in 2 nucleotide 3’ overhangs, characteristic of canonical Dicer-2 cleavage products (Fig. 3B,D). The average z-score across all three genome segments increased from 1.98 at 3 dpi to 2.54 at 6 dpi, suggesting sustained Dicer-2 activity throughout viral replication. Notably, virus-derived small RNAs in the 20-24 nucleotide range all displayed this complementarity pattern across the L, M, and S viral segments, indicating uniform processing.

**Fig. 3.**
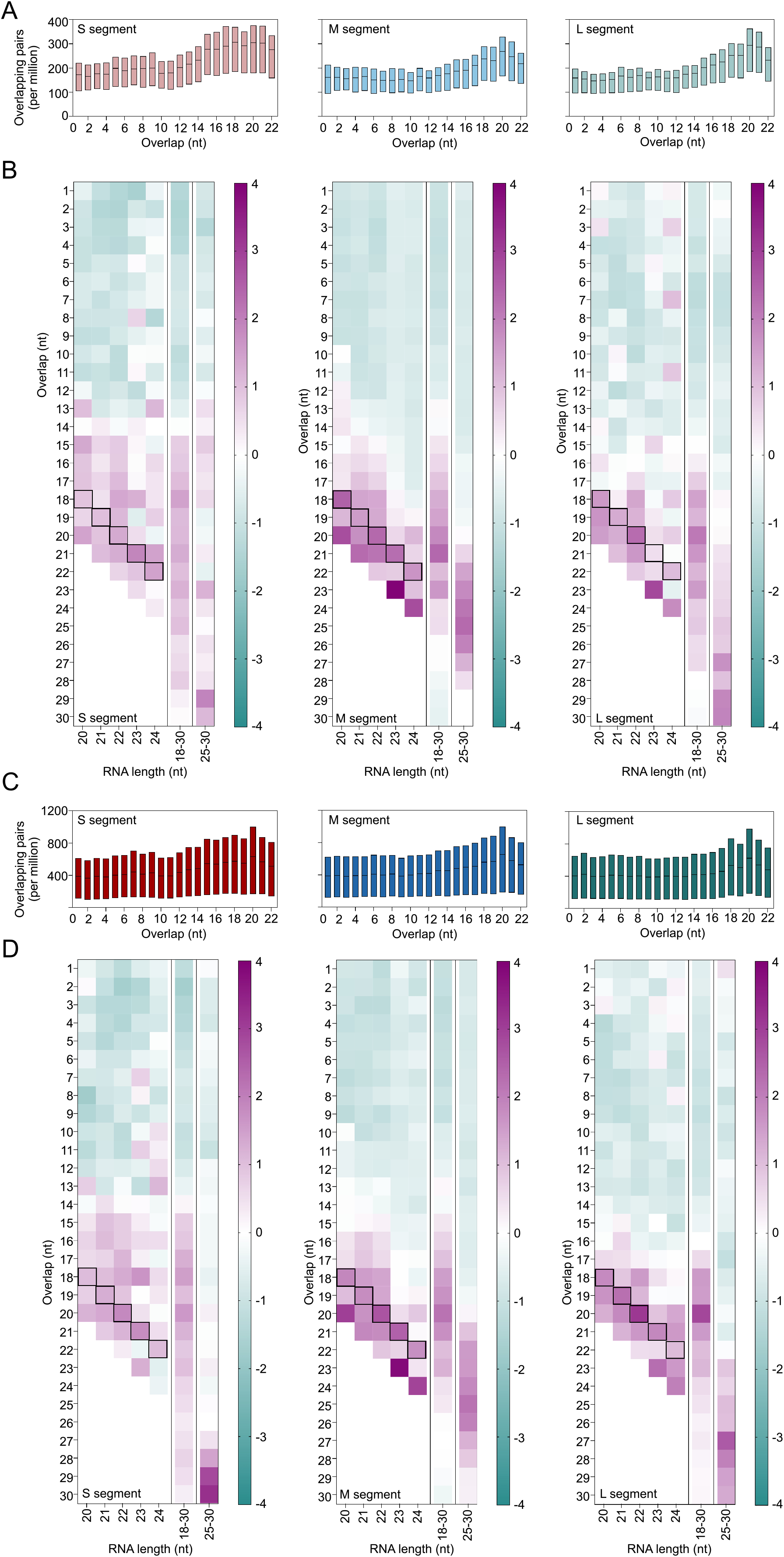
SFTSV virus-derived siRNAs exhibit the canonical 2-nt 3’ overhang signature of Dicer-2 processing. Reanalysis of data from Fig. 2. Dicer-2 processes double-stranded viral RNA into 20–22 nt duplexes with characteristic 2-nucleotide 3’ overhangs on each strand, a hallmark of antiviral RNA interference in arthropods. **(A**,**C)** Distribution of overlapping 22 nt vsiRNA pairs by nucleotide overlap length across S (left), M (middle), and L (right) genome segments (normalized per million mapped SFTSV reads). A peak at 2-nt overlap reflects precise Dicer-2 cleavage products. **(B**,**D)** Heat maps showing overlap probability z-scores for vsiRNAs of different lengths (x-axis) and overlap positions (y-axis) from three independent samples. Black boxes highlight the characteristic 2 nt overlap signature of Dicer-2 processing. Colour scale represents z-score values from −4 (teal) to +4 (purple). **(A**,**B)** data from 3 dpi. and **(C**,**D)** data from 6 dpi. The increase in overlap z-scores from 3 to 6 dpi indicates sustained Dicer-2 processing and amplification of antiviral vsiRNA production. Absence of 24–30 nt ‘ping-pong’ overlaps confirms that the tick antiviral response operates via the siRNA, not the piRNA, pathway.

No evidence of antiviral vpiRNAs (viral PIWI-interacting RNAs) was detected in the expected 24-30 nt range in infected tick cells at either timepoint. As a methodological control, SFV-infected AF525 cells displayed the characteristic sharp 19 nucleotide overlap signature resulting from impaired passenger strand processing in the absence of functional Ago2, along with piRNA signatures in the 24-30 nucleotide range with 10 nucleotide overlaps, as previously documented (11) (SI Appendix, Fig. S4). This contrasts with the broader overlap patterns observed in tick cells, confirming the host species-specific nature of antiviral RNA processing mechanisms.

### Viral RNAi suppressor partially rescues replication-defective SFTSV in tick cells

The findings above suggested that SFTSV NSs functions as an RNA silencing suppressor in tick cells. To test this hypothesis, we engineered a recombinant delNSs SFTSV expressing the tomato bushy stunt virus (TBSV) p19 protein, a well-characterized VSR that sequesters siRNAs to counteract antiviral RNA interference (38, 39). Both delNSs-p19V5 and delNSs-eGFP control viruses replicated efficiently in mammalian Vero E6 cells, confirming viral viability and demonstrating again that NSs is dispensable for replication in IFN-deficient mammalian systems (SI Appendix, Fig. S5).

In contrast, these two viruses showed markedly different phenotypes in tick cells. Wild-type SFTSV established robust infection with abundant viral N protein foci in BME/CTVM6 cells (Fig. 4A), while delNSs-eGFP showed no detectable infection by immunofluorescence (Fig. 4B). Functional complementation with p19 partially rescued this replication defect, enabling limited viral persistence with discrete N protein foci at 6 dpi (Fig. 4C). Quantification of immunofluorescent images demonstrated significantly more N protein-positive cells in delNSs-p19V5 versus delNSs-eGFP infected monolayers, indicating more efficient replication (Fig. 4D). Western blot analysis confirmed increased viral protein expression in delNSs-p19V5 infections, while minimal viral protein was detected in delNSs-eGFP lysates (Fig. 4E). Finally, growth kinetics revealed that delNSs-p19V5 achieved only intermediate replication levels with declining viral titers over time, demonstrating partial but incomplete functional rescue compared to wild-type virus (Fig. 4F). While this represented a modest non-significant improvement over delNSs-eGFP, both mutants remained severely attenuated. These complementation and quantification studies support a role for NSs in counteracting tick antiviral responses. To elucidate the underlying mechanism, we next investigated the biochemical properties of NSs.

**Fig. 4.**
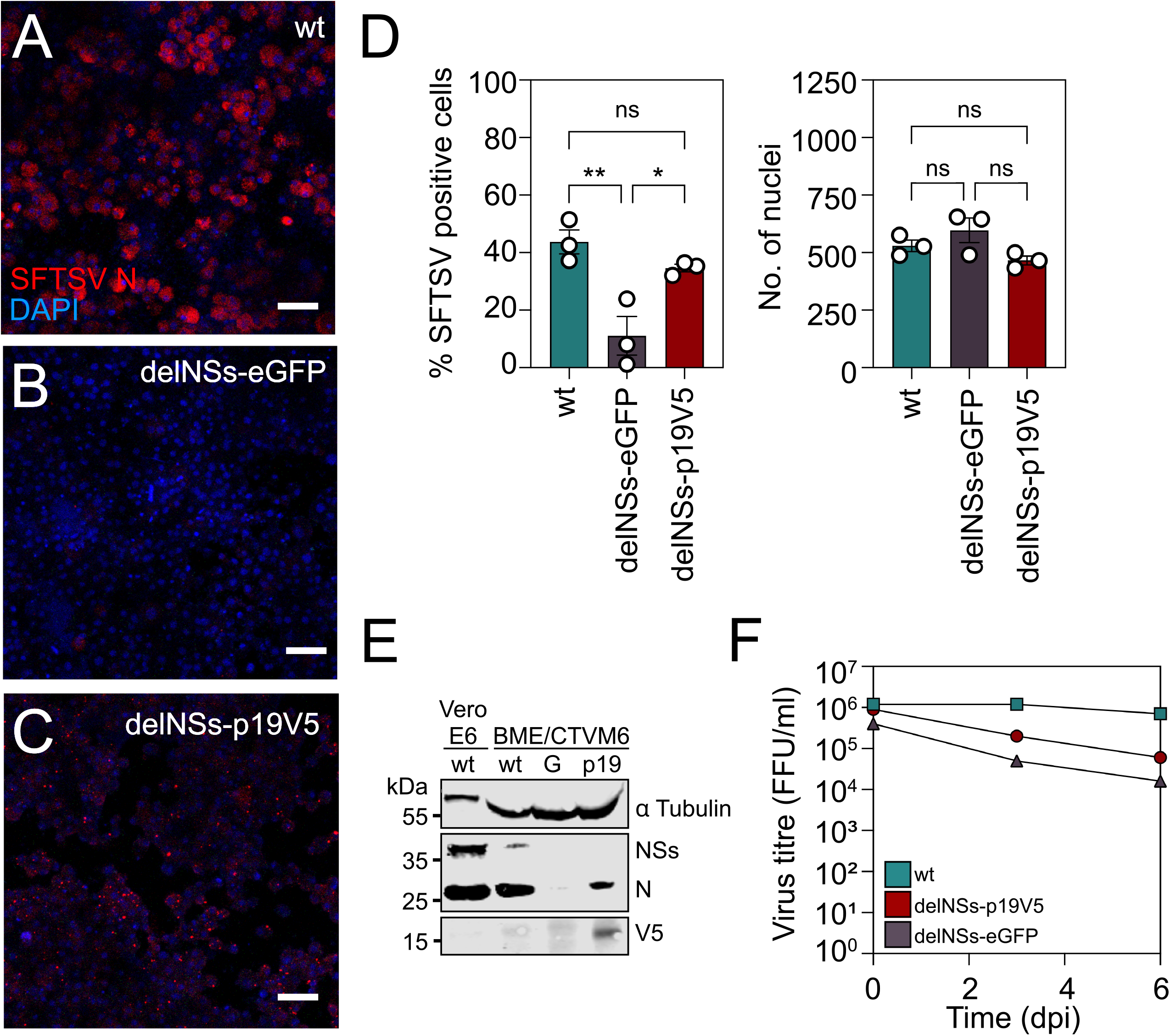
TBSV p19 protein rescues delNSs SFTSV replication in BME/CTVM6 cells. BME/CTVM6 cells infected with **(A)** wt SFTSV, **(B)** delNSs-eGFP or **(C)** delNSs-p19V5 virus (MOI: 10). At 6 dpi, cells were fixed and immunostained for SFTSV N protein (red) and nuclei (DAPI, blue). Imaging was carried out using a Zeiss LSM 880 Confocal Microscope, scale bar is equal to 50 μm. **(D)** (Left panel) Percentage of BME/CTVM6 cells infected with wt SFTSV (teal), delNSs-eGFP (red) or delNSs-p19V5 virus (purple) quantified from **(A-C)** at the timepoints indicated, evidenced by positive staining with anti-SFTSV N antibody. Data are plotted as the mean percent SFTSV-positive cells ± SD of n=3 randomised fields of view. (Right panel) Number of nuclei enumerated per field of view from left panel. Data are plotted as the mean number of nuclei observed ± SD of n=3 randomised fields of view. **(E)** Western blot analysis of viral protein expression in infected BME/CTVM6 cells (6 dpi, MOI: 10) and control Vero E6 cells (72 h p.i., MOI: 1). Blots were probed with antibodies against tubulin (loading control), SFTSV NSs, SFTSV N, or V5 tag. Lane labels: wt, wild-type SFTSV; G, delNSs-eGFP; p19, delNSs-p19V5. **(F)** Viral replication kinetics (MOI: 0.1) in BME/CTVM6 cells infected with wt SFTSV (teal), delNSs-p19V5 (red), or delNSs-eGFP (black) viruses. Virus titres were determined by focus-forming assay at indicated time points. Data show one representative experiment from n=3 independent repeats. Statistical analysis was performed using an ordinary one-way ANOVA, with Tukey’s multiple comparison test, with a single pooled variance. Asterisks indicate significance: * p ≤ 0.05, ** p ≤ 0.01; ns = not significant.

### SFTSV NSs exhibits structural features consistent with RNA-binding activity

Given that RNAi often requires direct interaction with RNA substrates, we investigated whether NSs functions as an RNA-binding protein. We generated a high-confidence structural model of the NSs protein using AlphaFold2 (40, 41) (Fig. 5A and SI Appendix, Fig. S6). The predicted structure revealed a compact globular protein with multiple α-helical regions and β-sheet domains. Electrostatic surface analysis demonstrated several distinct positively charged patches distributed across the protein surface (Fig. 5B), a characteristic feature of nucleic acid-binding proteins that facilitates interaction with the negatively charged phosphate backbone of RNA. We next employed computational approaches to identify potential RNA-binding domains (RBDs) within the NSs sequence. RBDetect analysis, which identifies RNA-binding sites based on sequence homology to known cellular RNA-binding sequences (42, 43), revealed two discrete regions with significant similarity to established RBDs: residues SER13-ARG22 and TRP133-THR141 (Fig. 5C; Dataset S1). The probability scores for RNA-binding activity peaked at approximately 0.9 for the first region and 0.6 for the second region, indicating strong predicted binding potential. To validate these predictions using an independent approach, we applied BindUP analysis, which predicts RNA-binding sites based on surface physicochemical properties (44) (Table S1). This analysis identified a region within α-helix 9 as having high probability for nucleic acid interaction. Notably, when the outputs from both RBDetect and BindUP were mapped onto the AlphaFold2 structural model, the predicted binding sites showed substantial overlap with spatial continuity (Fig. 5D), strengthening confidence in these predictions. Importantly, all identified putative RBDs were located within regions of high structural confidence in the AlphaFold2 model (SI Appendix, Fig. S6E).

**Fig. 5.**
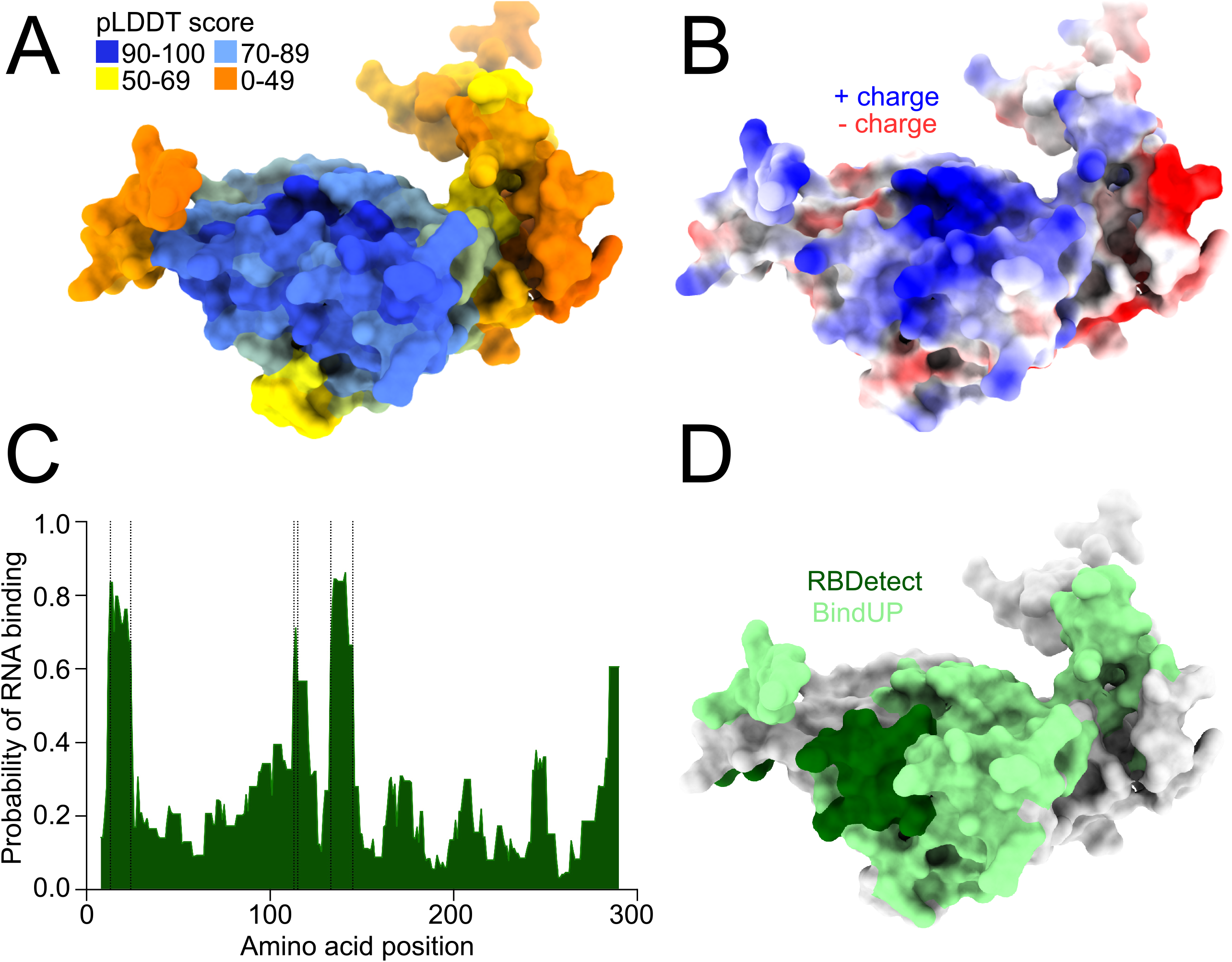
AlphaFold structural prediction reveals putative nucleic acid binding domains in SFTSV NSs protein. Predicted three-dimensional structure of SFTSV NSs protein (NCBI Protein: AJD86041). **(A)** Surface representation coloured by pLDDT confidence scores: very high confidence (dark blue, 90-100), confident (light blue, 70-89), low confidence (yellow, 50-69), very low confidence (orange, 0-49). **(B)** Electrostatic surface potential showing positive (blue) and negative (red) charges. **(C)** RNA-binding site prediction using RBDetect analysis. Plot shows probability of RNA binding (y-axis) versus amino acid position (x-axis). Regions between dotted lines indicate high predicted RNA-binding propensity. **(D)** Predicted RNA-binding residues mapped onto the protein surface: RBDetect predictions (dark green) and BindUP predictions (light green).

### SFTSV NSs binds single-stranded 22 nt vsiRNAs in infected cells

To validate NSs as an RNA-binding protein, we performed immunoprecipitation of V5-tagged p19 and NSs from recombinant virus-infected BME/CTVM6 cells at 2 dpi. Although protein expression levels in whole cell lysates varied between replicates, both proteins were detected in immunoprecipitated samples (Fig. 6A). Small RNA-seq analysis of co-immunoprecipitated RNAs revealed that p19 lysates were enriched for 19-22 nt RNAs as expected (45), whereas NSs immunoprecipitates showed enrichment for 22 nt RNAs (the characteristic products of Dicer-2 processing in tick cells) which comprised 50-55% of all detected reads across three independent replicates (Fig. 6B).

**Fig. 6.**
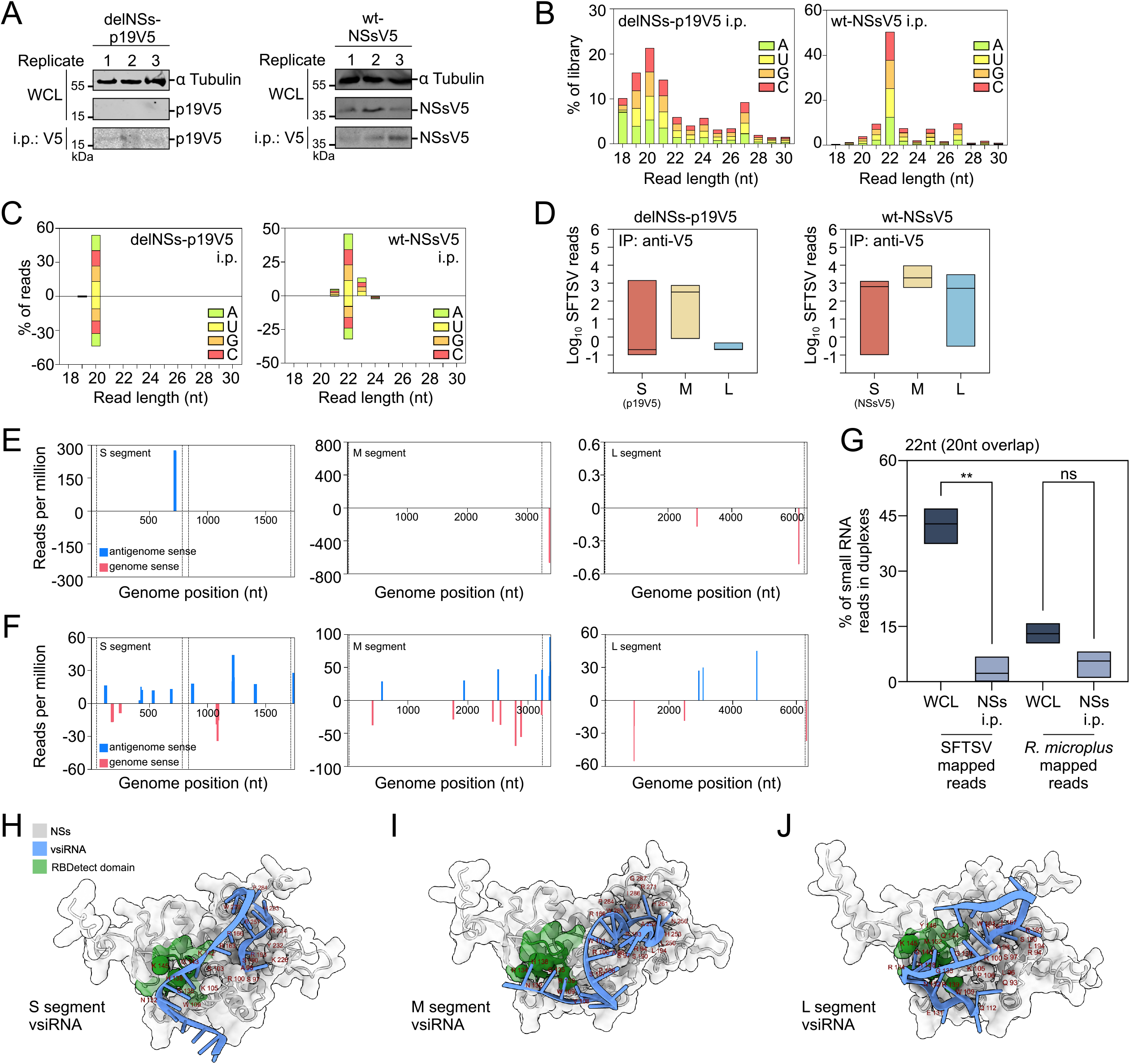
SFTSV NSs binds viral single-stranded RNA in infected BME/CTVM6 cells. **(A)** Immunoprecipitation of V5-tagged NSs proteins to assess NSs-vsiRNA interactions during SFTSV infection. BME/CTVM6 cells infected with SFTSV expressing either deletion mutant NSs-p19V5 (delNSs-p19V5) or wild-type V5-tagged NSs (wt-NSsV5) were harvested at 2 dpi and subjected to immunoprecipitation using anti-V5 antibody. Whole cell lysates (WCL; input) and immunoprecipitates (IP) were analysed by Western blot using antitubulin (loading control) and anti-V5 antibodies. RNA was subsequently extracted from immunoprecipitates for small RNA sequencing analysis. Molecular weights (kDa) are indicated. Data represent n=3 biological replicates (Replicate 1-3). **(B)** Histogram of all small RNA reads (18-30 nt) in immunoprecipitated samples, p19V5 i.p. (left panel) or NSsV5 i.p. (right panel) representative of n=3 biological replicates. **(C)** Small RNA reads from (B) mapped to SFTSV antigenomic (positive values) and genomic (negative values) strand RNAs from p19V5 i.p. (left panel) and NSsV5 i.p. (right panel) samples at 2 dpi. Read counts are coloured by the identity of the 5′ nucleotide and grouped by read length. Data are shown as the mean percentage of total mapped reads (y-axis); representative of n=3 biological replicates. **(D)** Distribution of small RNA reads (18-30 nt) mapping to SFTSV genome segments from p19V5 i.p. (left panel) and NSsV5 i.p. (right panel) samples at 2 dpi Data represent n=3 biological replicates. **(E)** Mapping of 18-25 nt SFTSV-derived vsiRNAs across the S (left panel), M (middle panel), and L (right panel) genome segments in p19V5 i.p. samples (reads per million). Data show mapping to the SFTSV antigenomic (blue) or genomic (pink) strands; numbers indicate nucleotide positions. Representative data from n=3 biological replicates. **(F)** Mapping of 22-nt SFTSV-derived vsiRNAs across genome segments in NSsV5 i.p. samples, as described in (C). Representative data from n=3 biological replicates. **(G)** Histogram showing the percentage of 22-nt small RNA reads with 20-nt complementary overlap mapping to SFTSV genome or *R. microplus* genome (GCA_002176555.1) in whole cell lysate (WCL) and NSsV5 i.p. samples. Data represent n=3 biological replicates. Statistical analysis was performed using Brown-Forsythe and Welch’s ANOVA with Dunnett’s T3 multiple comparison test to compare percentage of RNA reads in duplexes between samples. Asterisks indicate significance: * p ≤ 0.05, ** p ≤ 0.01; ns = not significant. **(H-J)** AlphaFold3-predicted structure of SFTSV non-structural protein NSs, shown as ribbon and surface representation (grey) bound to single-stranded viral small interfering RNA (vsiRNA, blue) generated during SFTSV infection of BME/CTVM6 tick cells (Fig. 6F) derived from the viral **(H)** S, **(I)** M and **(J)** L segments. Critical amino acids involved in RNA/protein binding to distances <4 Å are highlighted (brown). Green regions highlight key RNA-binding domains identified by RBDetect (Fig. 5C,D).

Analysis of small RNA composition demonstrated that 22 nt reads accounted for 75% of mapped reads derived primarily from SFTSV RNA in NSs immunoprecipitates, with approximately 45% from antigenomic sense and 30% from genomic sense sequences. In contrast, p19-associated SFTSV siRNAs were approximately 20nt in length, though this varied between replicates (Fig. 6C; SI Appendix, Fig. S7). When comparing normalized read counts across viral genome segments, both proteins showed similar binding levels, although siRNAs derived from the SFTSV M segment were those most reproducibly immunoprecipitated by NSs across replicates (Fig. 6D).

Genome mapping analysis revealed distinct siRNA mapping patterns between the two proteins. The p19-associated 18-25 nt siRNAs mapped to limited viral genome regions (Fig. 6E, SI Appendix, Fig. S8), whereas NSs-bound 22 nt RNAs mapped to multiple reproducible hotspots distributed across all three viral segments (Fig. 6F, SI Appendix, Fig. S9). Notably, hotspots derived from antigenomic sense S segment RNA corresponded to regions within the genome with strongly predicted secondary structures, such as the siRNA hotspot identified mapping to nt position 1210-1238 of the antigenomic S RNA, which corresponds to the head region of a hairpin structure (Fig. 6F; SI Appendix, Fig. S10).

Crucially, NSs immunoprecipitation revealed a statistically significant reduction (**p ≤ 0.01) in the proportion of 22 nt siRNA duplex pairs (those with 20 nt overlaps), decreasing from approximately 45% in total infected cell lysates to less than 10% in NSs-bound samples. This reduction was specific to vsiRNAs, as *R. microplus* genome-derived siRNAs showed no difference in duplex abundance (Fig. 6G). Size-specificity analysis confirmed similar single-strand enrichment for 21 nt and 23 nt siRNAs but not for 24 nt or 28 nt species, demonstrating that NSs selectively binds vsiRNA single strands within a defined size range (SI Appendix, Fig. S11). This sequestration is consistent with post-Dicer processing of vsiRNAs, as correctly processed vsiRNA duplexes were robustly and reproducibly detected in infected tick cells (Fig. 3). These findings support a model in which NSs suppresses RNAi through selective engagement and depletion of single-stranded 22-nt vsiRNAs. The observed reduction in duplex vsiRNA abundance, combined with structural predictions, provides strong support for this mechanism. However, direct evidence of impaired Argonaute loading, such as through Ago2 immunoprecipitation or silencing reporter assays, has yet to be demonstrated. How NSs binding modulates vsiRNA availability to influence RNAi outcomes, thereby permitting productive viral replication, remains to be determined. The molecular details and stoichiometric requirements of NSs–vsiRNA interactions warrant further investigation.

To validate our computational predictions, we performed molecular modelling using authentic 22-nucleotide vsiRNAs identified from NSs immunoprecipitation (Table S2). Docking analyses revealed multiple RNA-binding interfaces within 4Å contact distance for vsiRNAs derived from all three SFTSV genomic segments: S (Fig. 6H), M (Fig. 6I), and L (Fig. 6J).

Notably, the predicted RNA contact residues clustered within the same protein region identified by both RBdetect and BindUP algorithms (Fig. 5D), providing highly convergent evidence for the location of functional RNA-binding sites from orthogonal computational approaches. This structural validation strongly supports the siRNA-binding capacity of SFTSV NSs protein in tick cells and establishes a molecular framework for understanding NSs-mediated RNAi suppression.

To further interrogate the predicted RNA-binding interface of SFTSV NSs, we computationally generated a panel of charge-reversal mutants targeting conserved basic residues within and adjacent to the predicted binding pocket (Table 1; Arg100, Lys105, Arg139, and Lys145). Initial in silico modelling across independent computational approaches, including AlphaFold3-based complex prediction, HADDOCK docking, and iPTM interface confidence analysis, consistently localised RNA interaction to a region spanning residues 100-145 (more strictly between residues 135-144), independent of RNA segment, indicating a stable and well-defined predicted binding site (Fig. 5 and Fig. 6H–J). Confidence in the predicted folding of the mutant NSs structures remained high, indicating there was no overall loss of structure mediated by the mutations (SI Appendix, Fig. S12A). Superposition of the in silico–generated structural models of wild-type NSs with these mutants revealed no substantial changes to the overall fold or local backbone conformation, with the mutated residues occupying equivalent spatial positions in all models. This suggests that the introduced substitutions are not predicted to induce gross structural rearrangement, but rather alter local interface chemistry (SI Appendix, Fig. S12B-E).

**Table 1.**
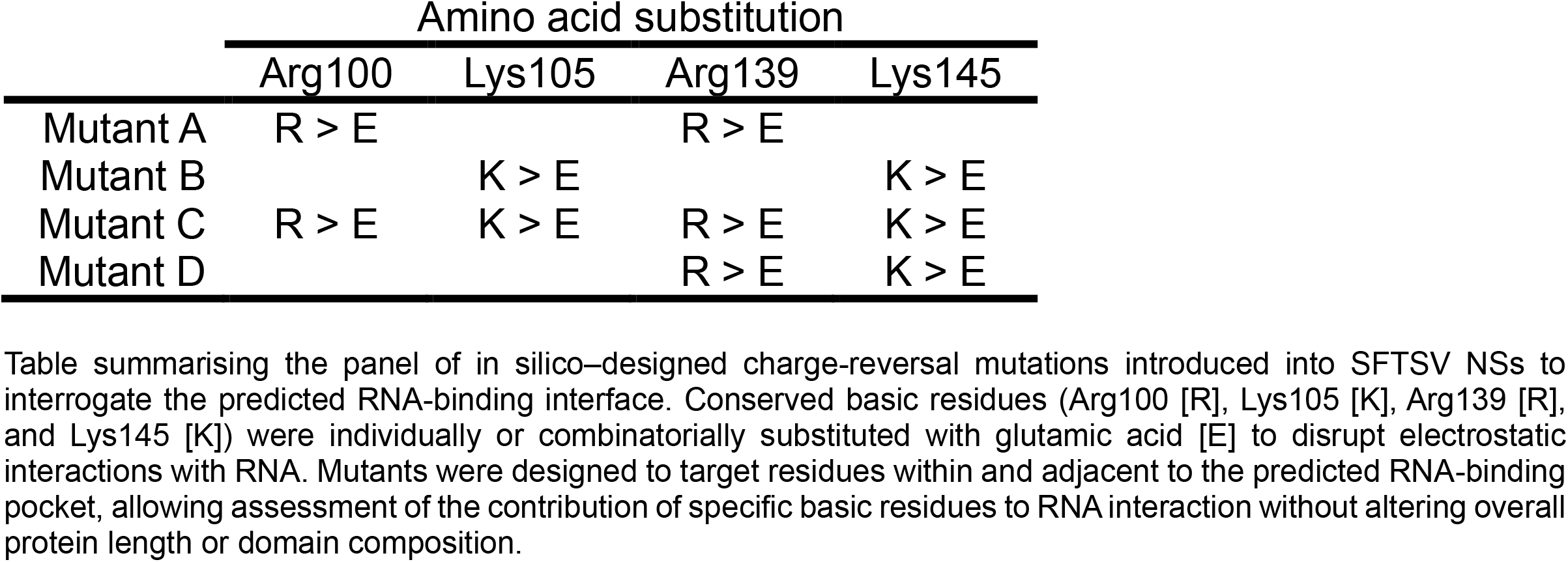
Design of SFTSV NSs RNA-binding interface mutants. Table summarising the panel of in silico–designed charge-reversal mutations introduced into SFTSV NSs to interrogate the predicted RNA-binding interface. Conserved basic residues (Arg100 [R], Lys105 [K], Arg139 [R], and Lys145 [K]) were individually or combinatorially substituted with glutamic acid [E] to disrupt electrostatic interactions with RNA. Mutants were designed to target residues within and adjacent to the predicted RNA-binding pocket, allowing assessment of the contribution of specific basic residues to RNA interaction without altering overall protein length or domain composition.

We observed reduced protein/RNA interactions particularly with NSs Mutant C and Mutant D, suggesting weakened or less well-defined protein/RNA interfaces (Fig. 7A). We therefore selected these to assess the impact of these mutations on predicted RNA binding using single-stranded vsiRNAs derived from the S (RNA_3), M (RNA_5), and L (RNA_10) genome segments. Across all RNAs, charge-reversal mutations resulted in reduced predicted binding scores and a displacement of the interaction interface away from the mutated residues, as reflected by concordant reductions in interface confidence (Fig. 7A) and decreases in the predicted interface area and binding free energy across both AlphaFold3 and HADDOCK models (Fig. 7B). Mutations targeting residues within the core 133–145 region had the strongest effects, with Mutant D (Arg139Glu, Lys145Glu) showing the most pronounced and consistent reduction in predicted binding, particularly for M- and L-segment-derived RNAs. Together, these analyses support a model in which NSs engages viral RNAs through a defined, charge-dependent interface centred on residues 133–145, rather than through nonspecific electrostatic interactions.

**Fig. 7.**
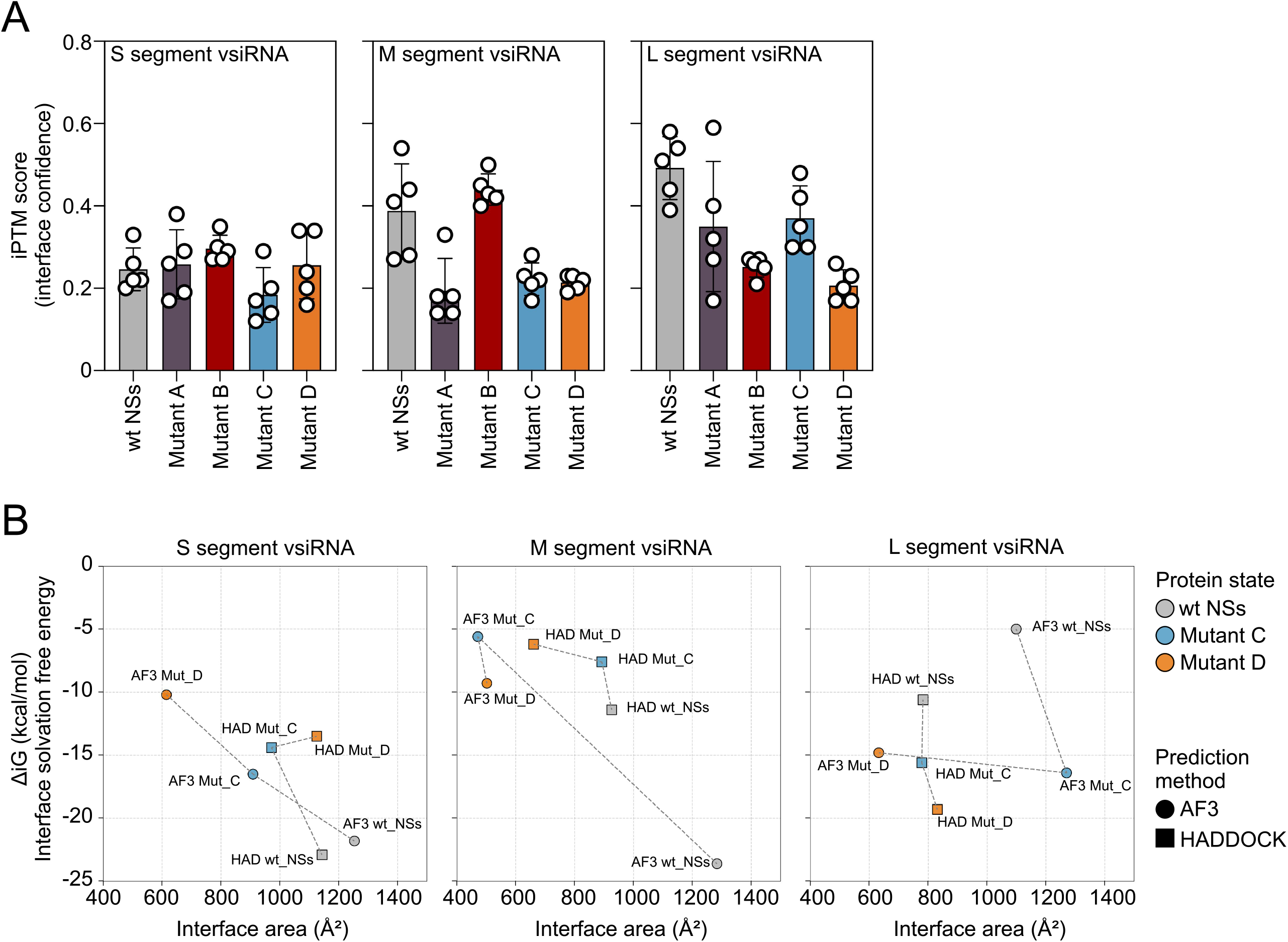
Effect of protein mutations on viral siRNA binding predicted through structural modelling. **(A)** Mean interface predicted TM-score (iPTM) values for interactions between the wild-type protein (wt NSs) or mutant variants (Mutant A–D) and vsiRNAs derived from the S, M, or L genome segments. Bars represent the mean iPTM score calculated from five AlphaFold3 models per complex, with error bars indicating standard deviation. Individual data points show iPTM scores from each AlphaFold3 model. **(B)** Relationship between protein–RNA interface area and interface solvation free energy (ΔiG) calculated using PDB ePISA for wild-type and mutant complexes. Circles denote AlphaFold3-derived complexes (AF3) and squares denote HADDOCK-derived complexes (HAD). Colours indicate protein variants (wt NSs, grey; Mutant A, purple; Mutant B, red; Mutant C, blue; Mutant D, orange). Dashed lines connect AlphaFold and HADDOCK predictions for the same protein–RNA pair. Reduced iPTM scores, smaller interface areas, and less favourable ΔiG values in mutant complexes indicate weakened protein–RNA binding compared with the wild-type protein.

## DISCUSSION

This study reveals a fundamental dichotomy in SFTSV replication strategies between mammalian and arthropod hosts, establishing NSs as a critical viral RNAi suppressor essential for tick cell infection. The striking observation that NSs protein is virtually undetectable in productively-infected tick cells, despite being absolutely required for viral replication, challenges conventional paradigms of viral protein function and highlights the sophisticated evolutionary adaptations that enable arboviral transmission.

The species-specific permissivity patterns observed across tick cell lines (SI Appendix, Fig. S1A,B) provide crucial insights into possible SFTSV vector competence. The successful replication in *R. microplus* cells (BME/CTVM6 and BME/CTVM23) correlates with the known epidemiological distribution of SFTSV-positive ticks, suggesting that cellular factors governing viral entry and replication are phylogenetically conserved traits that determine vector competence in nature. Cell lines from two other species, *Rhipicephalus decoloratus* (BDE/CTVM16) and *Hyalomma anatolicum* (HAE/CTVM9) originating from East Africa and India respectively, also supported SFTSV replication, although the virus has not been reported from these tick species or geographic regions, suggesting the potential for spread beyond its currently known distribution. The sustained viral production over extended culture periods (SI Appendix, Fig. S1C,D) confirms the establishment of infections expected of arthropod-borne bunyaviruses (25, 28, 46, 47). The failure of SFTSV to replicate in *Ixodes* spp. cell lines, despite these ticks vectoring related phenuiviruses (48), indicates that host range determinants may operate at the cellular level.

Here we show that SFTSV NSs protein expression is repressed in tick cells yet is unexpectedly essential for sustained virus replication. This contrasts with mosquito-borne bunyaviruses such as Rift Valley fever virus, where NSs repression does not impact virus replication in mosquito cells and instead facilitates viral persistence (47, 49–51), suggesting distinct host-specific requirements for NSs in different arthropod vectors. The paradoxical requirement for NSs in tick cells represents a remarkable evolutionary balancing act that extends beyond recent findings in mammalian systems (23). Our demonstration that delNSs SFTSV fails to sustain infectious progeny in tick cells (Fig. 1B) and showed no detectable nucleocapsid protein (Fig. 1C), contrasts sharply with dispensable NSs function in IFN-deficient mammalian cells (34). Proteomic analysis revealed differential temporal expression of SFTSV proteins in tick cells, with structural proteins (N, GPC) and the RNA-dependent RNA polymerase (RdRp) showing significant increases from 3 to 6 dpi, while the non-structural protein NSs remained at consistently low levels throughout infection. The detection of NSs by mass spectrometry despite its absence in Western blots suggests tight regulation of this protein in infected tick cells, perhaps compounded by antibody sensitivity (Fig. 1I,J). The paradox of NSs essentiality despite undetectable protein levels may be resolved through several non-exclusive mechanisms. Firstly, based on established proteomics intensity-to-concentration relationships (52, 53), the observed mean peptide intensity for SFTSV NSs (Log2 = 15) suggests protein concentrations in the low nanomolar range (1-10 nM) range, which, while below Western blot detection limits, align with the nanomolar binding affinities characteristic of other viral RNA silencing suppressors (54). While our results support a vsiRNA depletion model, NSs activity is most consistent with interference at or around the RISC engagement stage; however, we cannot yet resolve whether this occurs through effects on vsiRNAs prior to Ago2 loading or through interactions with single-stranded guide RNAs and/or Ago2-containing RISC complexes after loading. Secondly, NSs may exhibit exceptionally high RNA-binding affinity or cooperative binding that amplifies its suppressive capacity at low concentrations. Thirdly, temporal or subcellular compartmentalization may concentrate NSs activity during critical early infection phases, when RNAi activation would otherwise prevent productive replication. The profound replication defect observed in NSs-deficient infections (Fig. 1E,F) reinforces the notion that even low-abundance NSs is functionally essential in tick cells. These mechanisms may collectively explain how limited quantities of NSs achieve effective RNAi suppression in the tick host.

The selective reduction of NSs subgenomic mRNA in tick cells provides a mechanistic explanation for the protein’s low abundance while maintaining its essential function. Our Northern blot analysis (Fig. 1L) demonstrates that while both genomic and antigenomic viral RNAs accumulate to similar levels in tick and mammalian cells, NSs mRNA shows markedly reduced levels specifically in tick cells, suggesting a post-transcriptional regulation mechanism unique to arthropod hosts. This regulatory strategy represents an evolutionary compromise that maximizes NSs activity during critical early infection phases while minimizing detection by host surveillance mechanisms.

The demonstration that SFTSV infection triggers robust small RNA production in tick cells (Fig. 2A) establishes RNAi as a primary antiviral defence mechanism. Our beta elimination analyses provide definitive evidence that the observed 22-nucleotide small RNAs represent authentic virus-derived siRNAs rather than degradation products (Fig. 2B-E). This finding significantly extends previous work by Xu *et al*., who demonstrated vsiRNA production during SFTSV infection of whole *H. longicornis* ticks through blood feeding (22). Our study advances this understanding by providing detailed mechanistic insights into vsiRNA processing, demonstrating temporal dynamics of the RNAi response, and establishing the functional consequences for viral replication in cell culture systems.

The emergence of methylated vsiRNAs at 6 dpi (Fig. 2E), which are absent at 3 dpi, supports a model wherein the host RNAi machinery eventually overcomes transient viral suppression. Given that NSs is expressed at low levels in infected tick cells (undetectable by Western blot; Fig. 1C), complete sequestration of all vsiRNAs would present a significant stoichiometric challenge. This constraint is further amplified by the temporal dynamics of antiviral RNAi responses, as vsiRNA accumulation increases substantially between 3 and 6 dpi. The appearance of RISC-loaded vsiRNAs at 6 dpi therefore indicates that sustained RNAi pathway activation can ultimately saturate the suppressive capacity of this low-abundance viral suppressor, enabling functional antiviral silencing despite initial evasion mechanisms.

Our observations support a ‘depletion strategy’ whereby viral suppressors of RNAi achieve functional suppression without requiring abundant expression. To address potential stoichiometric limitations, we propose that NSs operates through high-affinity binding kinetics rather than simple mass action. Assuming dissociation constants comparable to characterised VSRs [Kd ~0.1-1 nM (54)] and estimated cellular vsiRNA concentrations of 10-100 nM during peak infection, even low-abundance NSs could plausibly compete for substrate binding. Interference with vsiRNAs during RISC engagement, rather than competition with fully assembled RISC complexes, would be expected to be more efficient, providing a conceptual explanation for how limited NSs quantities could achieve functional suppression despite abundant total vsiRNA pools. Such high-affinity interactions are characteristic of other VSRs, including tombusvirus p19, which functions at substoichiometric concentrations (Kd ~0.17 nM) and achieves substantial siRNA depletion despite limited protein abundance (45).

The progressive amplification of vsiRNA accumulation across all viral segments between 3- and 6-dpi (Fig. 2F-H) indicates that tick cells mount increasingly robust antiviral responses over time. The concentrated targeting within the unencapsidated S segment intergenic region (55) and NSs open reading frame provides direct mechanistic evidence for RNAi-mediated NSs mRNA degradation, explaining the reduced NSs protein levels observed. This segment-specific targeting pattern suggests that viral RNA secondary structures or transcript abundance influence RNAi machinery accessibility, with implications for understanding viral evolutionary constraints.

The demonstration of canonical Dicer-2-mediated vsiRNA production through strand overlap analysis (Fig. 3) confirms that tick antiviral RNAi pathways follow established arthropod paradigms. The predominance of 20-22 nucleotide products with characteristic 2-nucleotide 3’ overhangs indicates uniform processing across all viral segments (8). Notably, the absence of detectable vpiRNA responses contrasts with mosquito antiviral systems (46, 56–59), suggesting lineage-specific adaptations in arthropod small RNA-mediated immunity. However, the complete absence of vpiRNA production warrants further mechanistic investigation to determine whether this reflects genuine pathway absence or differential regulation.

The successful complementation of NSs-deficient virus with the heterologous p19 suppressor (Fig. 4) provides strong evidence that NSs functions as a VSR in tick cells. The partially restored viral protein expression and intermediate replication kinetics in delNSs-p19V5 infected cells demonstrate functional complementation while revealing that NSs possesses unique properties beyond simple siRNA sequestration. This incomplete complementation suggests NSs may: (i) exhibit specific binding preferences for SFTSV-derived vsiRNAs through evolved sequence or structural recognition motifs, (ii) interact with additional host factors beyond the core RNAi machinery, or (iii) function through temporal coordination with other viral proteins during the replication cycle. The heterologous p19 suppressor, while functionally competent, lacks these virus-specific adaptations that have co-evolved to optimise NSs performance within the SFTSV life cycle.

Our integrated structural and functional analyses establish NSs as a novel siRNA-binding protein that selectively engages single-stranded 22 nt vsiRNAs through size-specific molecular recognition. This finding extends observations from plant-infecting bunyaviruses, particularly Tospoviruses, where NSs proteins bind both double-stranded siRNAs and miRNAs with varying specificity depending on phylogenetic origin; American clade tospoviruses exhibit broad-spectrum binding, whereas Eurasian clade NSs proteins demonstrate more restricted binding to short dsRNA substrates (20, 60, 61). In these plant-infecting systems, NSs suppresses RNA silencing by binding and sequestering small interfering RNAs, thereby preventing their incorporation into the RNA-induced silencing complex (20, 61, 62). While these observations derive from plant host systems, they suggest that RNA binding-mediated suppression of RNAi may represent a conserved strategy employed by bunyaviral NSs proteins across diverse hosts. Our findings extend this concept to tick cells and support a model in which SFTSV NSs similarly limits antiviral RNAi through interaction with virus-derived small RNAs. Notably, while RNAi suppressor activity has not been clearly demonstrated for NSs proteins from vertebrate-infecting bunyaviruses in either vertebrate hosts or arthropod vectors (20), our results provide strong mechanistic evidence for this function.

Earlier studies implicated SFTSV NSs in modulation of RNA interference primarily through interactions with Argonaute proteins, based on protein–protein interaction data generated in mammalian cells (23) or indirect vsiRNA profiling from whole feeding ticks (22). In contrast, our study directly interrogates NSs function in tick-derived cells and demonstrates association of NSs with single-stranded 22-nt vsiRNAs in the relevant host context. These findings shift the mechanistic focus from Argonaute-centric models toward vsiRNA engagement as a central feature of NSs-mediated RNAi suppression in ticks, providing a plausible explanation for why previous Ago2 co-immunoprecipitation studies detected vsiRNA-associated complexes without resolving functional Argonaute loading. While we cannot exclude additional NSs–Ago2 interactions, our data support a model in which vsiRNA sequestration contributes substantially to RNAi suppression in tick cells. Together, this work defines a tick-specific framework for NSs function that is distinct from mechanisms inferred in mammalian systems and refines our understanding of how SFTSV adapts to the arthropod RNAi pathway.

While our immunoprecipitation and computational modeling provide strong evidence for NSs-vsiRNA interactions, quantitative binding parameters remain to be determined. The inherent challenges associated with NSs biochemical characterization, including insolubility during bacterial expression (24, 63) and extensive C-terminal disorder that complicates *in vitro* folding, have limited direct binding studies. Despite these technical constraints, our cellular interaction data and structural predictions provide compelling concordant evidence for the RNA-binding capacity and size-specific vsiRNA recognition observed in infected tick cells.

The convergence of computational predictions, AlphaFold2 structural modelling (Fig. 5), and experimental data strongly supports a model in which NSs engages virus-derived siRNAs through a defined RNA-binding interface, with discrete binding hotspots suggesting recognition of specific viral RNA secondary structural features (Fig. 6, SI Appendix, Fig. S10). Together with the enrichment of single-stranded 22-nt vsiRNAs in NSs immunoprecipitates, these findings support a model in which NSs suppresses antiviral RNAi through direct engagement of functional vsiRNAs in tick cells. This interaction is most consistent with interference at or around the stage of RISC engagement following Dicer processing, although the precise temporal relationship to Ago2 loading remains to be determined. Future studies employing Ago2 immunoprecipitation or functional RNAi reporter assays will be required to resolve this final mechanistic detail.

Importantly, our overlap analysis coupled with molecular docking experiments provide strong evidence for the single-stranded nature of the bound RNA substrates, identifying a previously unrecognised mode of RNAi suppression among bunyaviruses that diverges from canonical dsRNA-binding suppressors. The molecular docking validation using authentic vsiRNA sequences derived from viral infections provides direct structural confirmation and establishes a comprehensive molecular framework for understanding NSs-mediated RNAi evasion. This mechanistic insight represents a substantial functional advancement beyond the previous protein-protein interaction studies (NSs/Ago2) conducted in mammalian cell systems.

The robustness of the proposed mechanism is further supported by the convergence of multiple independent *in silico* analyses, all of which point to a discrete and functionally relevant RNA-binding interface within NSs (Fig. 7, SI Appendix, Fig. S12). The selective sensitivity of RNA binding to charge-reversal mutations argues against nonspecific electrostatic association or global protein destabilisation, and instead supports engagement through a defined, charge-dependent interaction surface. The harmony between distinct computational frameworks, together with functional data obtained in arthropod cells, provides supportive evidence for interface-specific RNA recognition by SFTSV NSs.

Our findings highlight a key evolutionary adaptation in tick-borne phenuiviruses: the requirement for RNAi suppression in arthropod vectors, but not mammalian hosts. This host-specific dependency likely constrains viral evolution and presents an opportunity for vector-targeted antiviral strategies. The identification of NSs as a siRNA-sequestering protein, rather than an upstream pathway inhibitor, reveals a novel mechanism of immune evasion and offers potential targets for future intervention strategies. Disrupting NSs/vsiRNA interactions or enhancing guide RNA stability could block viral replication in ticks without affecting mammalian infection. While our findings are derived from established tick cell line models, *in vivo* validation in whole *H. longicornis* or *R. microplus* spp. ticks will be essential to confirm the physiological relevance of NSs-mediated RNAi suppression during natural infection. These insights establish NSs as a critical determinant of SFTSV vector competence and uncover a previously unrecognised function for this viral protein.

## MATERIALS AND METHODS

Tick- and mammalian-cell infection experiments were performed using recombinant SFTSV generated by reverse genetics. Viral replication was assessed by immunofocus assays, immunofluorescence microscopy, Western blotting, Northern blotting and proteomics. Small RNA responses were analysed using methylation-sensitive small RNA sequencing, immunoprecipitation of V5-tagged viral proteins, and computational mapping pipelines. Structural predictions and protein–RNA docking were performed using AlphaFold2/3, HADDOCK, and complementary RNA-binding prediction algorithms. Full experimental procedures and computational pipelines are provided in the SI Appendix.

## ACKNOWLEDGEMENTS

We thank Prof. Ulrike Munderloh, University of Minnesota, for permission to use the ISE6, AAE2 and AAE12 cell lines. We also thank Dr Matthew Arnold (Royal Veterinary College) for support with the initial AlphaFold modelling.

## AUTHORS’ CONTRIBUTIONS

Conceptualization: M.F., M.McF., E.S., A.C., A.K., B.B., Data Curation: M.F., M.McF., R.H.P., R.A., A.T.C., M.P., B.B., Formal Analysis: M.F., M.McF., R.H.P., R.A., A.T.C., W.K., R.D.S., U.S.L., M.P., A.C., A.K., B.B., Funding Acquisition: B.B., Investigation: M.F., M.McF., R.H.P., R.A., A.T.C., W.K., K.D., R.D.S., U.S.L., A.C., A.K., B.B., Methodology: M.F., M.McF., R.H.P., R.D.S., U.S.L., M.P., E.S., B.B., Project administration: B.B., Resources: L.B.S., B.B., Software: R.H.P., W.K., M.P., B.B., Supervision: U.S.L., A.C., A.K., B.B., Validation: M.F., M.McF., R.H.P., R.A., A.T.C., W.K., K.D., R.D.S., U.S.L., M.P., E.S., A.C., A.K., B.B., Visualization: M.F., M.McF., R.H.P., R.A., A.T.C., W.K., B.B., Writing – original draft: M.F., M.McF., A.K., B.B., Writing – review & editing: M.F., M.McF., R.H.P., R.A., A.T.C., W.K., K.D., R.D.S., U.S.L., L.B.S., M.P., E.S., A.C., A.K., B.B.

## DATA AVAILABILITY

All data that support the findings of this study are openly available from Enlighten Research Data (https://doi.org/10.5525/gla.researchdata.2028). All sequencing data generated in this study have been deposited in the NCBI Sequence Read Archive under BioProject accession number PRJNA1283909. The bioinformatic scripts used for analysis can be found in the Zenodo online repository (https://doi.org/10.5281/zenodo.15877198).

