## Supporting Appendix for "SFTSV NSs protein is a novel tick antiviral RNAi response suppressor"

### Supporting Information Text

#### MATERIALS AND METHODS

##### Experimental model and subject detail

**Cell culture**

Vero E6 cells (1) were cultured in DMEM (Gibco) supplemented with 10% foetal bovine serum (FBS) (Gibco). HuH7-Lunet-T7 cells(2), which stably express T7 RNA polymerase were obtained from R. Bartenschlager, Heidelberg University and were grown in DMEM (Gibco) supplemented with 2 mM L-glutamine, nonessential amino acids, and 10% FBS (Gibco). All mammalian cells were cultured at 37 °C with 5% $CO_2$. The tick cell lines used in this study were *Amblyomma americanum* AAE2 (3) and AAE12 (4), *Amblyomma variegatum* AVL/CTVM17(5), *R. microplus* BME/CTVM2(5), BME/CTVM6(5), and BME/CTVM23(6), *R. decoloratus* BDE/CTVM16(5), *H. anatolicum* HAE/CTVM9(7), *Ixodes scapularis* ISE6(8),*; Ixodes ricinus* IRE/CTVM19 and IRE/CTVM20 (9) and *Rhipicephalus sanguineus* RML-RSE(10, 11), sourced from the Tick Cell Biobank at the University of Liverpool. *The tick cell lines were grown at either 28 °C or 32 °C in sealed flat-sided culture tubes (Nunc) containing 2 ml of appropriate complete culture medium* (L-15, L-15B, L-15B300, L-15/MEM, L-15/H-Lac, L-15/L-15B or L-15/H-lac/L-15B) (5, 12, 13)as described previously *for maintenance,* and transferred to sealed T25 or T80 non-vented flasks to bulk for experimentation. Aag2-AF525 cells, an Ago2 knockout cell line derived from Aag2-AF5 (*Aedes aegypti* mosquito; European Collection of Cell Cultures (ECACC): 19022601) cells by CRISPR/Cas9 (14, 15) were maintained in Leibovitz's L-15 medium supplemented with 10% FBS (Gibco), 10% tryptose phosphate broth (Gibco), and penicillin-streptomycin (final concentration 100 units/ml, 100 µg/ml; Gibco) and maintained at 28 °C.

**Plasmids**

Plasmids for the recovery of SFTSV have been described previously(16, 17). The expression plasmids pTM1-HB29L and pTM1-HB29N contain the SFTSV HB29 L and N ORFs under control of the T7 promoter and encephalomyocarditis virus internal ribosome entry site sequence. The rescue plasmids pTVT7-HB29S, pTVT7-HB29M, and pTVT7-HB29L contain full-length cDNAs of the SFTSV strain HB29 antigenome segments flanked by T7 promoter and hepatitis delta ribozyme sequences. Plasmid pTVT7-HB29SNSsV5 encodes an S segment expressing NSs with a C-terminal V5 epitope tag (GKPIPNPLLGLDST). Plasmid pTVT7-HB29SdelNSs:TBSVp19V5 encodes an S segment in which the NSs ORF is replaced with the tomato bushy stunt virus p19 protein, also with a C-terminal V5 epitope tag (TBSV; Accession No. M21958.1).

**Generation of recombinant viruses from cDNA**

Recombinant SFTSV were generated by transfecting HuH7-Lunet-T7 cells with 0.1 µg pTM1-HB29L, 0.5 µg pTM1-HB29N, and 1 µg each of pTVT7-based plasmids expressing wt or recombinant viral antigenomic segments. Transfections used 3 µl TransIT-LT1 (Mirus Bio LLC) per µg of DNA as the transfection reagent. After 5 days, transfection supernatants were collected, clarified by low-speed centrifugation, and stored at -80°C. Recombinant virus stocks were amplified in Vero E6 cells at 37 °C by infection at an MOI of 0.01, with culture medium harvested at 7 dpi. The genome segments of recovered viruses were amplified by RT-PCR and sequenced to confirm that no mutations had occurred during the rescue process.

**Virus Production**

SFTS viruses used in this study (rHB29pp [referred to as 'wt SFTSV'] (17)or recombinant viruses rHB2912aaNSs [delNSs](16), rHB29delNSsGFP [delNSs eGFP](16), rHB29NSsV5 [NSsV5] and rHB29delNSsp19V5 [p19V5]) were generated using reverse genetics technologies. The Hubei 29 (HB29) strain is based on a plaque-purified stock called Hubei 29pp (HB29pp) provided by Amy Lambert (CDC Arbovirus Diseases Branch, Fort Collins, CO). Working stocks of SFTSV were generated in Vero E6 cells by infecting at a low multiplicity of infection and harvesting the cell culture medium 7 dpi. Recombinant virus stocks were confirmed by Sanger sequencing to ensure no mutations had occurred during propagation. Previous data have demonstrated that recombinant viruses have the same replication kinetics and viral protein expression profiles as the wt/parental viruses in time course of infection studies(16, 17).

**Virus titration by immunofocus assays**

Virus titres were determined by focus-forming assays in Vero E6 cells. Confluent Vero E6 cells were infected with 10-fold serial dilutions of virus prepared in phosphate-buffered saline containing 2% FBS and incubated for 1 h at 37 °C. Following infection, cells were overlaid with MEM supplemented with 2% FBS and 0.6% Avicel (FMC Biopolymer). Cells were incubated for 6 days before fixation and immunostaining as described previously(17).

#### RNA extraction from cell cultures

Total RNA was extracted from cultured cells using 1 ml TRIzol reagent (Thermo Fisher Scientific) according to the manufacturer's protocol. RNA concentrations were measured using a NanoDrop spectrophotometer, and RNA was stored at -80 °C until use.

#### Western blotting

Cell lysates were prepared at indicated timepoints by adding 300 µl lysis buffer (100 mM Tris-HCl pH 6.8, 4% SDS, 20% glycerol, 200 mM DTT, 0.2% bromophenol blue, and 25 U/ml Pierce Universal Nuclease [Thermo Fisher]). Proteins were separated on 4-12% SDS-polyacrylamide gradient gels (Invitrogen) and transferred to Hybond-C Extra membranes (Amersham). Membranes were blocked in saturation buffer (PBS containing 5% dry milk and 0.1% Tween 20) for 1 h at room temperature. Primary antibodies included SFTSV anti-N or anti-NSs polyclonal rabbit antibodies(17), anti-tubulin monoclonal antibody (Sigma), or anti-V5 antibody (Abcam). After washing, membranes were incubated with goat anti-rabbit DyLight 680 or goat anti-mouse DyLight 800 secondary antibodies (Thermo

Fisher). Proteins were visualized using a Li-Cor Odyssey CLx infrared imaging system. Data shown in the manuscript are representative gels from 2-3 independent replicates.

##### Immunofluorescence microscopy

BME/CTVM6 cells previously seeded onto 13mm coverslips for 24 h were infected with wt or recombinant SFTS viruses at indicated MOIs. At defined timepoints post-infection, cells were fixed with 8% paraformaldehyde in PBS for 15 min at room temperature, permeabilized with 0.5% Triton X-100 for 10 min and blocked with 4% milk in PBS for 1 h. Cells were incubated with primary antibodies overnight at 4 °C, washed 3x with PBS, followed by staining with Alexa Fluor-conjugated secondary antibodies for 1 h at room temperature. Nuclei were counterstained with DAPI (1 µg/ml). Images were acquired using a Zeiss LSM 880 confocal microscope and number of nuclei and infection rates quantified using ImageJ (version 1.52p).

##### Northern blotting

BME/CTVM6 or Vero E6 cells were infected with wt SFTSV, and total cellular RNA was extracted at 72 h post-infection using TRIzol reagent (Invitrogen). RNA samples (1 µg) were electrophoresed through 1.2% agarose gels in TAE buffer (18) and transferred via capillary action to positively charged nylon membranes (Roche). Membranes were hybridized with digoxigenin-labelled RNA probes (150 ng each) complementary to negative-sense genomic RNA, positive-sense antigenomic RNA, or subgenomic mRNA sequences encoding N or NSs proteins. Detection was performed using a DIG Northern Starter Kit (Roche) according to the manufacturer's instructions. Data shown in the manuscript are representative gels from 2-3 independent replicates.

##### Small RNA sequencing and library preparation

Total RNA (1 µg per sample) was extracted and processed using the BGI Genomics Small RNA Library Construction Protocol for DNBSEQ platform (DNBSeq, UMI small library, SE50). Following sequential 3’ and 5’ adaptor ligation, reverse transcription, and PCR amplification (16–17 cycles), libraries were PAGE-purified, circularized to single-stranded DNA, and quality-assessed using an Agilent Bioanalyzer. Sequencing was performed on a DNBSEQ-G400 instrument with 50 bp single-end reads.

##### Small RNA methylation analysis

Methylation status of small RNAs was determined using a beta-elimination assay as previously described(19). BME/CTVM6 cells (2.5 x $10^5$ cells/$cm^2$) were infected with wt SFTSV (MOI: 0.1) or mock-infected for 3 or 6 days. AF525 cells infected with SFV (MOI: 10, 24 h) served as positive controls for 2'-O-methylation. Following TRIzol LS extraction, RNA samples (200 µl) were treated with 5 µl 20x borate buffer (pH 8.6) and either 12.5 µl 200 mM sodium periodate or nuclease-free water (controls), incubated for 15 min at room temperature, then treated with

10 µl glycerol for an additional 15 min. RNA was precipitated overnight at -20 °C using GlycoBlue coprecipitant, 3 M sodium acetate (pH 5.2), and absolute ethanol, recovered by centrifugation (16,000 x g, 15 min, 4 °C), washed with 70% ethanol, and resuspended in 100 µL sodium borate buffer (55 mM, pH 9.5). RNA was purified using the Monarch RNA Cleanup Kit and sequenced by BGI Tech Solutions using DNBSEQ UMI technology (minimum 500 ng input, 20 million clean reads per sample).

#### Small RNA immunoprecipitation assay and RNA extraction

BME/CTVM6 cells were infected with either NSsV5- or p19V5- expressing SFTS viruses (MOI: 1) and cell lysates prepared at 2 dpi to assess viral protein-RNA interactions. Immunoprecipitation was performed using Protein G Dynabeads (Thermo Fisher). Lysis buffer contained 20 mM Tris-HCl pH 7.5, 150 mM NaCl, 5 mM $MgCl_2$, 0.5% NP-40, and protease/phosphatase inhibitors (Roche). Wash buffer consisted of 50 mM Tris-HCl pH 7.5, 200 mM NaCl, 1 mM EDTA, and 1% NP-40. Dynabeads (20 µL) were washed three times, conjugated with 5 µg anti-V5 antibody (Abcam) for 1 h at 4°C, then washed again. Cells were lysed in 100 µL lysis buffer per $10^6$ cells for 20 min on ice, clarified by centrifugation (16,000 x g, 20 min), and immunoprecipitated overnight at 4 °C with rotation. After three washes, proteins were eluted in LDS buffer at 90 °C for 10 min. For RNA extraction, eluates were treated with proteinase K (1 µg, 1 h, 50 °C; Sigma) then processed with TRIzol following standard phenol-chloroform extraction. RNA was precipitated with isopropanol, washed with 75% ethanol, and resuspended in nuclease-free water.

#### Small RNA computational analysis

##### Small RNA bioinformatics pipeline

Small RNA libraries (Table S3 and Table S4) were sequenced and processed using a comprehensive bioinformatics pipeline described on the Zenodo repository. Raw sequencing reads were quality-filtered and adapter-trimmed using fastp v0.23 (20, 21) (**1_small_RNA_fastp_processor.sh**) with small RNA-optimized parameters: length filtering (18-32 nt), Phred quality score threshold Q20, unqualified base percentage limit 40%, maximum 2 N bases per read, tail trimming with window size 1, poly-G and poly-X trimming enabled, and overrepresentation analysis activated. Adapter sequences were auto detected. FASTA files were standardized and formatted for optimal analysis (**2_fasta_formatter.sh**).

##### Read alignment and quality control

Reference genome indices were constructed using Bowtie v1.3.1 (22, 23) (**3_bowtie_index_builder.sh**), and trimmed reads were aligned to viral (SFTSV HB29 strain) and host (*R. microplus* GCA_002176555.1) reference genomes (**4_recursive_bowtie_alignment_pipeline_outputs_standspecificBAM.sh, 4a_bowtie_stringent_smallRNA_aligner.sh**) with stringent parameters: zero mismatches allowed (-v 0), maximum 5 mapping locations per read (-m 5), best alignment reporting (--best --strata), and exhaustive search (--tryhard). Postalignment filtering retained only uniquely mapping reads with MAPQ ≥20, excluding unmapped (SAM flag 4), secondary (flag 256), and supplementary (flag 2048) alignments. BAM files were strand-separated based on SAM flag 16 (reverse complement) using SAMtools v1.15(24). Quality filtering statistics and mapping efficiencies were tracked throughout the pipeline to ensure data integrity.

**Small RNA sequence analysis and characterization**

Read length distributions and nucleotide composition analysis were performed using custom scripts (**5_Length_Distribution_Analysis.sh**) and (**5a_strand_specific_Length_Distribution_Analysis.sh**) calculating first- and last-base frequencies (A, T/U, G, C percentages) across read lengths 1-50 nt individually, with reads >50 nt grouped. Data transformation for visualization was conducted using R scripts (**5_for_generating_length_vs_nt_pc_graph_data.R, 5a_for_generating_strand_specific_length_vs_nt_pc_graph_data.R**).

Chromosome-specific read counting was performed across all BAM files (**6_bam_chromosome_read_counter.sh**).

Coverage analysis employed a custom Python pipeline (**7_rpm_normalized_coverage_tool.py**) utilizing pysam v0.19.1(25) with CIGAR-aware position counting (equivalent to bedtools -split behaviour) and RPM normalization based on total input reads from original FASTQ files rather than mapped reads. The most abundant small RNA sequences were identified and characterized (**9_top_mapping_small_rna_finder.py**).

**Ping-pong amplification analysis**

Ping-pong amplification signatures (for vpiRNAs) were detected using a custom algorithm (**8_pingpong_smallRNA_analyzer.py**) analysing overlapping antisense small RNA pairs within a 1-19 nucleotide window, requiring complementary reads of 23-29 nt length with ≥20 nt overlap on opposite strands. The algorithm calculated Z-scores for overlap frequencies and identified statistically significant ping-pong pairs using both raw counts and probability-based H-signatures. Data consolidation across samples was performed (**8a_ping_pong_data_consolidator.R**) and formatted for analysis (**8b_ping_pong_matrix_formatter.R**).

**Small RNA classification**

Comprehensive small RNA characterization was conducted using an integrated pipeline (**10_small_rna_analysis.py**) with configuration management (**10a_small_rna_config.yaml, 10b_setup_and_run.sh**) that classified small RNA populations as miRNA-like (20-24 nt), siRNA-like (20-25 nt), or piRNA-like (24-32 nt) based on size distribution profiles. All analyses maintained strand-specific resolution with forward strand values reported as positive and reverse strand as negative for visualization.

#### AlphaFold prediction of SFTSV NSs

The sequence of SFTSV NSs (Hubei-29; GenBank ID: AJD86041) was used to query CollabFold's AlphaFold2 structure prediction algorithm using the advanced beta notebook(26, 27). Structure prediction was performed using MMseqs2 for multiple sequence alignment generation, with a maximum MSA depth of 512:1024 sequences and MSA subsampling enabled. Five models were generated with two random seeds each, using 3 recycle iterations and pTM scoring enabled. The ten resulting models were ranked by pLDDT scores, with the highest-ranking model (rank_1_model_5_ptm_seed_0; pLDDT: 72.77, pTMscore: 0.7104) selected for further structural analysis (SI Appendix, Fig. S5). Subsequent structural analysis was performed using UCSF ChimeraX(28).

#### SFTSV NSs RNA binding prediction

RNA binding prediction for the SFTSV NSs protein sequence was performed using RBDetect (29), a machine learning model trained by Shrinkage Discriminant Analysis (SDA) with a dataset of 8,891 experimentally identified polypeptides from RBDmap experiments, using positive examples (RNA-bound polypeptides) and negative examples (RNA-released polypeptides)(30). For each amino acid position in the NSs protein sequence, RBDetect assigns a probability value for RNA binding based on the fragment centred at that position. A Hidden Markov model was then used to visualize the probabilities sequentially, determining the most probable RNA binding regions across the protein structure. Predicted RNA binding sites (Dataset S1) were mapped onto the AlphaFold-predicted NSs structure to identify potential functional domains. We also utilised BindUP(31), a computational tool that predicts RNA-binding regions in protein sequences using physicochemical and structural features rather than sequence homology. The tool exploits characteristic surface properties of RNA-binding proteins (RBPs), including electrostatic potential, hydrophobicity, and shape complementarity, to distinguish them from non-RBPs. BindUP employs a machine-learning framework trained on curated RBP and non-RBP datasets. The software computes surface patches on protein structures (experimental or predicted) and scores their likelihood of mediating RNA interactions (31) (Table S1).

#### RNA secondary structure prediction

RNA secondary structures were predicted using the forna web server (http://rna.tbi.univie.ac.at/forna/, accessed May 2025) powered by the ViennaRNA package(32). The 76-nucleotide RNA sequence derived from nucleotides 1191-1266 of the antigenomic HB29 S RNA (GenBank: KP202165) was subjected to minimum free energy folding calculations using Turner energy parameters at 37°C. The predicted secondary structure with the lowest free energy was selected for analysis and visualization. No folding constraints were applied during the prediction process.

#### Modelling of RNA binding to SFTSV NSs

Structural modelling of SFTSV NSs-vsiRNA interactions was performed using AlphaFold3. The SFTSV NSs protein sequence (Hubei-29; GenBank ID: AJD86041) was submitted to the AlphaFold3 webserver (version 2.0) with representative 22-nucleotide vsiRNA sequences identified from NSs immunoprecipitation experiments in SFTSV-infected BME/CTVM6 tick cells. vsiRNA sequences were selected based on abundance in NSs-IP samples from identified hotspots within each viral RNA segment (Table S2). Five independent structural predictions were generated for each NSs-vsiRNA complex using default parameters. Models were selected based on per-residue confidence scores (pLDDT >70 for NSs functional domains) and interface confidence scores (>0.5 for protein-RNA contacts). Structural analysis was performed using UCSF ChimeraX (version 1.6)(28), focusing on RNA-binding interfaces, contact residues within 4 Å of vsiRNA substrates, and structural determinants of vsiRNA recognition (Dataset S2).

#### Protein-RNA complex predictions

Protein–RNA complex structures were predicted using AlphaFold3 (33) and HADDOCK (34) for the wild-type protein (wt NSs) and mutants (Table 1; mutant A–D). For AlphaFold predictions, five models per complex were generated. Mean iPTM scores and standard deviations were calculated to assess interface confidence and model consistency (mean ± SD iPTM scores). Protein–RNA interfaces generated through Alphafold3 and HADDOCK were analysed using PDB ePISA(35), to extract either Interface area (Å²) or Interface solvation free energy (ΔiG, kcal/mol). Results were visualised as ΔiG versus interface area scatter plots, comparing prediction methods and protein variants.

#### Proteomic analysis

To estimate relative abundance of SFTSV proteins, we analyzed proteomic data from project PXD054068, which comprises whole-cell proteomes of BME/CTVM6 tick cells infected with SFTSV (MOI 1) at 3- and 6-dpi with four biological replicates. Corresponding peptides for each viral protein [non-structural protein (NSs; AJD86041), nucleocapsid protein (N; AJD86040), glycoprotein precursor (GPC; AJD86039) or RNA-dependent RNA polymerase (RdRp; AJD86038)] were pooled across replicates at each time point, and peptide intensity distributions were analyzed. Scripts used to generate plots and statistics are available from the Zenodo online repository (https://doi.org/10.5281/zenodo.15877198). Statistical significance of changes in peptide intensities between time points was assessed using Welch's t-test.

#### Biosafety

All experiments with infectious SFTSV were performed in the Richard Elliott Biosafety Laboratories (REBL) at the MRC-University of Glasgow Centre for Virus Research, under containment level 3 (CL-3) conditions, approved by the UK Health and Safety Executive (GM223/20.2a).

#### Quantification and Statistical Analysis

Experiments were performed with a minimum of n=2 biological replicates unless otherwise stated, with each biological sample tested in technical replicates where applicable. Technical replicates were averaged before statistical analysis. Data are presented as mean ± SD e.g. for virus titres and cell counts. For immunofluorescence quantification, n=3 randomised fields of view were analysed per condition. Small RNA sequencing experiments used n=3 biologically independent samples. Statistical significance for multiple group comparisons was assessed using Brown-Forsythe and Welch's ANOVA with Dunnett's T3 multiple comparison test or an ordinary one-way ANOVA with Tukey's multiple comparisons test, assuming a single pooled variance. Statistical significance was set at $p \leq 0.05$, with * $p \leq 0.05$, ** $p \leq 0.01$; 'ns' indicates not significant. Individual sample sizes are indicated in figure legends. Statistical analyses were performed using GraphPad Prism Version 10.5.0(673).

### Figures

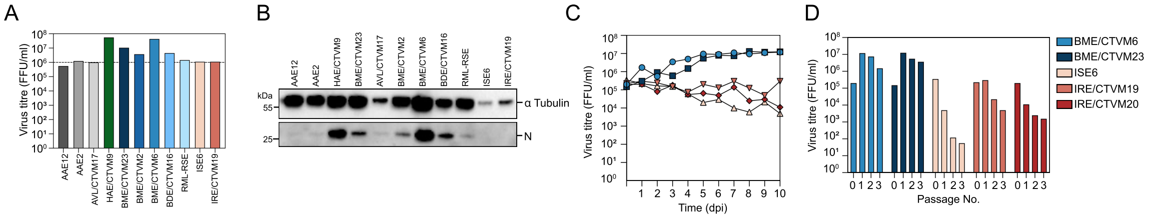


**Fig. S1. Initial characterisation of wt SFTSV replication in a diverse range of tick cells.**
**(A)** Titre of wt SFTSV in the supernatant of infected tick cells (MOI: 1) assayed at 10 dpi. Cell lines used were derived from tick genera *Amblyomma* (grey), *Hyalomma* (purple), *Rhipicephalus* (blue) or *Ixodes* (pink). Dashed line represents titre of initial inoculum. Data are plotted as virus titre (FFU/ml) and show a representative experiment of n=2 biological replicates.
**(B)** Representative images of cell extracts derived from cell monolayers in **(A)**. Western blots were probed with anti-tubulin (loading control) or anti-SFTSV N antibodies to detect cellular and viral protein expression.
**(C)** Tick cell lines derived from *Rhipicephalus microplus* or *Ixodes* spp. were infected with wt SFTSV (MOI: 0.1) for up to 10 dpi. Data are plotted as virus titre (FFU/ml) and show n=1 experiment.
**(D)** At 10 dpi, wt SFTSV-infected tick cell cultures described in **(C)** were passaged and split (1:2) and left to recover for a minimum of 14 days. This process was repeated for up to 3 subsequent passages. Data are plotted as virus titre (FFU/ml) and show a representative experiment (n=1).

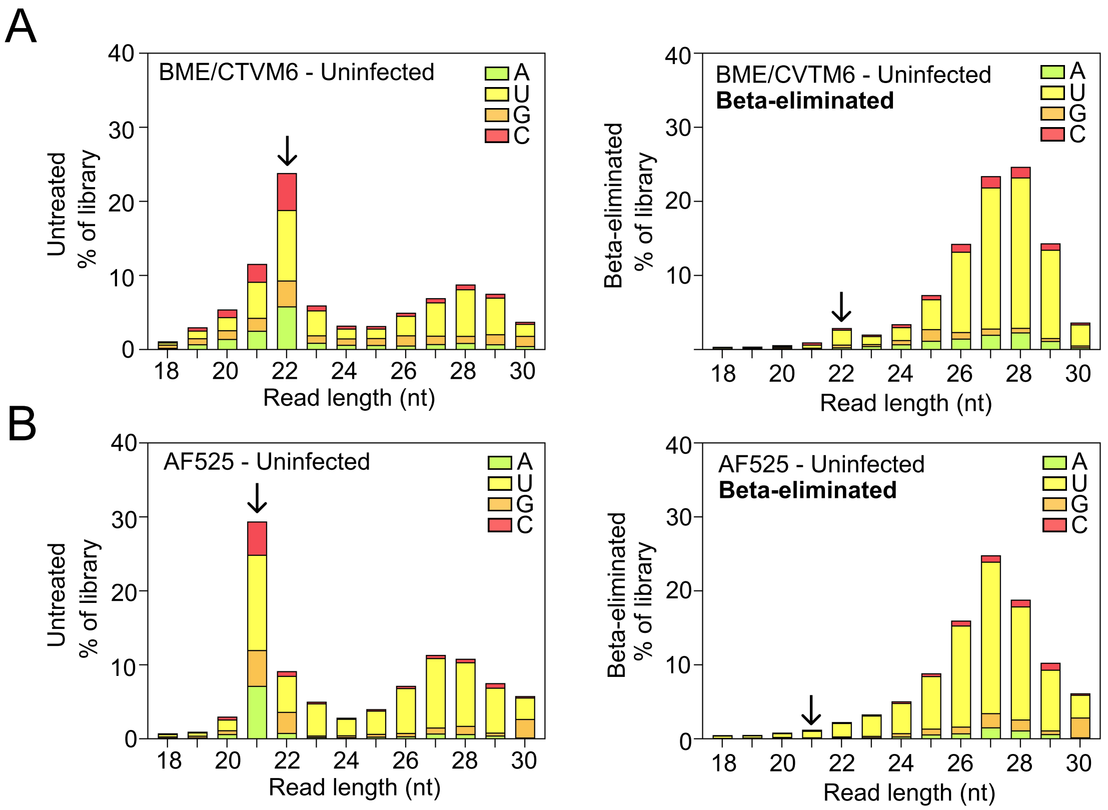


**Fig. S2. Beta-elimination on uninfected tick and mosquito cell cultures.**
Histogram of all small RNA reads (18-30 nt) from **(A)** uninfected BME/CTVM6 or **(B)** AF525 (Ago2 knock out) cells after 3 days in culture. RNA was extracted from cell cultures and samples were either untreated (left hand panel) or beta-eliminated (right hand panel). Arrows indicate the siRNA species of RNA depleted in beta-eliminated conditions. Read counts are coloured by the identity of the 5′ nucleotide and grouped by read length. Data are shown as mean reads as a percentage of the total small RNA sequencing library (y-axis); representative of n=3 biologically independent samples.

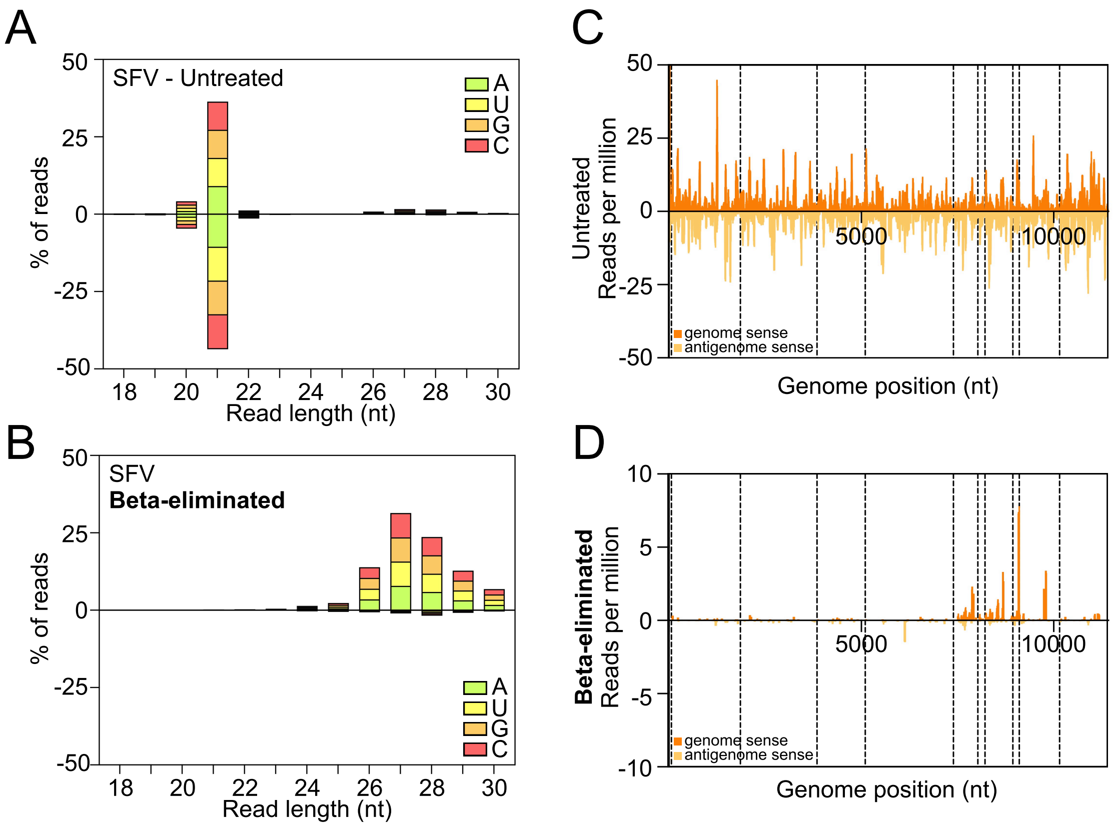


**Fig. S3. Beta-elimination in Semliki Forest virus-infected mosquito cell cultures.**
An *Aedes aegypti* Ago2 knock out mosquito cell line (AF525) was infected with SFV (MOI: 10). At 24 h p.i., total cell RNA was extracted and either left untreated (top) or subjected to beta-elimination (bottom). Histograms of small RNA reads (18-30 nt) from **(A)** untreated or **(B)** beta-eliminated samples that mapped to the SFV genomic (negative values) and antigenomic (positive values) strand RNAs in infected cells. Read counts are coloured by the identity of the 5′ nucleotide and grouped by read length. Data are shown as the mean percentage of total mapped reads (y-axis); representative of n=3 biologically independent samples. **(C, D)** SFV-derived 21 nt vsiRNAs in **(C)** untreated or **(D)** beta-eliminated samples. Coverage across the viral genome for genomic (orange) and antigenomic (yellow) sense RNAs is shown (y-axis: vsiRNA reads per million) representatives of n=3 biologically independent samples.

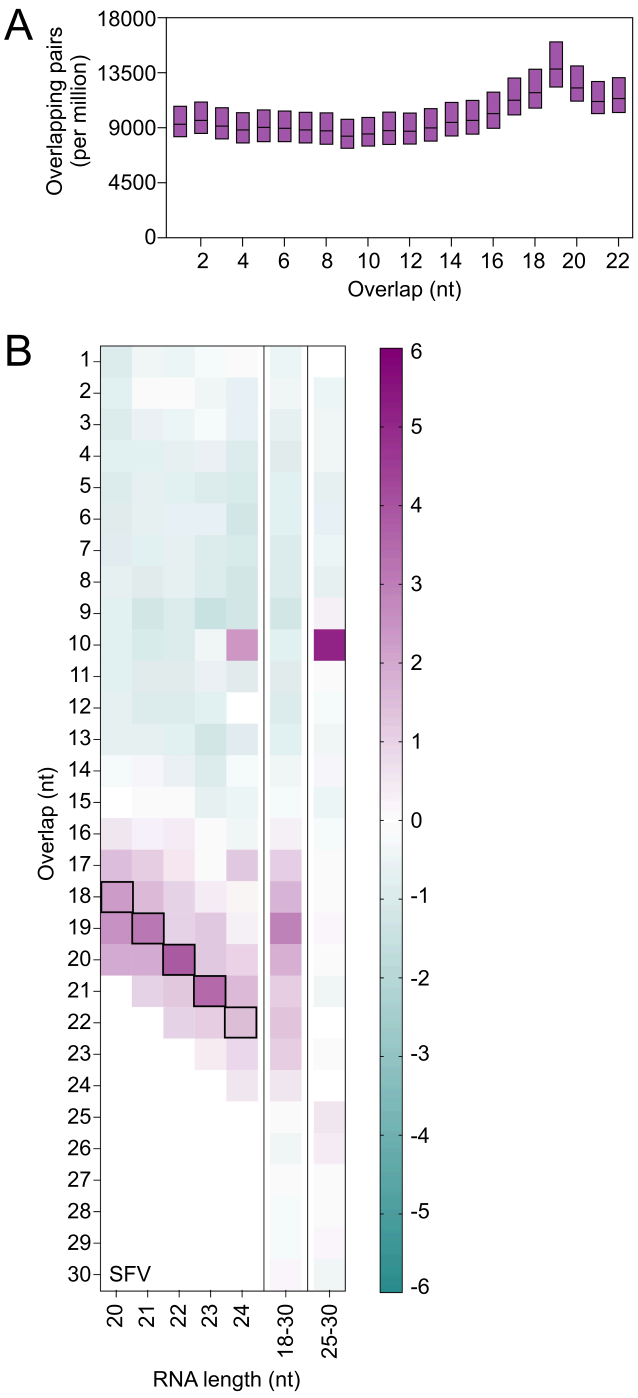


**Fig. S4. Distribution of overlapping 21 nt vsiRNA pairs in SFV-infected AF525 cells.**
Reanalysis of data from SI Appendix, Fig. S3. **(A)** Distribution of overlapping 21 nt vsiRNA pairs by nucleotide overlap length across the SFV genome (SFV4, GenBank ID: KP699763), presented as normalised per million mapped SFV reads. **(B)** Heat maps showing overlap probability z-scores for vsiRNAs of different lengths (x-axis) and overlap positions (y-axis) from three independent samples. Black boxes highlight the characteristic 2 nt overlap signature of Dicer-2 processing. Colour scale represents z-score values from -6 (teal) to +6 (purple).

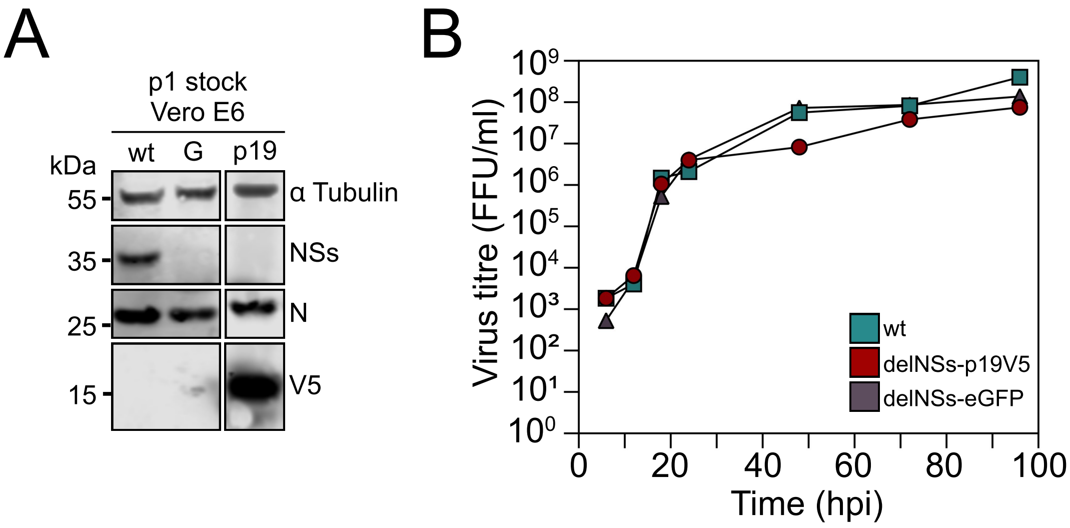


**Fig. S5. Characterisation of delNSs-p19V5 SFTSV in Vero E6 cells.**

**(A)** Western blot analysis of viral protein expression in infected Vero E6 cells (6 dpi; MOI: 1). Blots were probed with antibodies against tubulin (loading control), SFTSV NSs, SFTSV N, or V5 tag. Lane labels: wt, wild-type SFTSV; G, delNSs-eGFP; p19, delNSs-p19V5. Viral replication kinetics in Vero E6 cells infected with wild-type SFTSV (teal), delNSs-p19V5 (red), or delNSs-eGFP (purple) viruses. Virus titres were determined by focus-forming assay at indicated time points. Data show one representative experiment from n=3 independent experiments.

**(B)** Viral replication kinetics in Vero E6 cells infected with wild-type SFTSV (teal), delNSs-p19V5 (red), or delNSs-eGFP (black) viruses. Virus titres were determined by focus-forming assay at indicated time points. Data show one representative experiment from n=3 independent experiments.

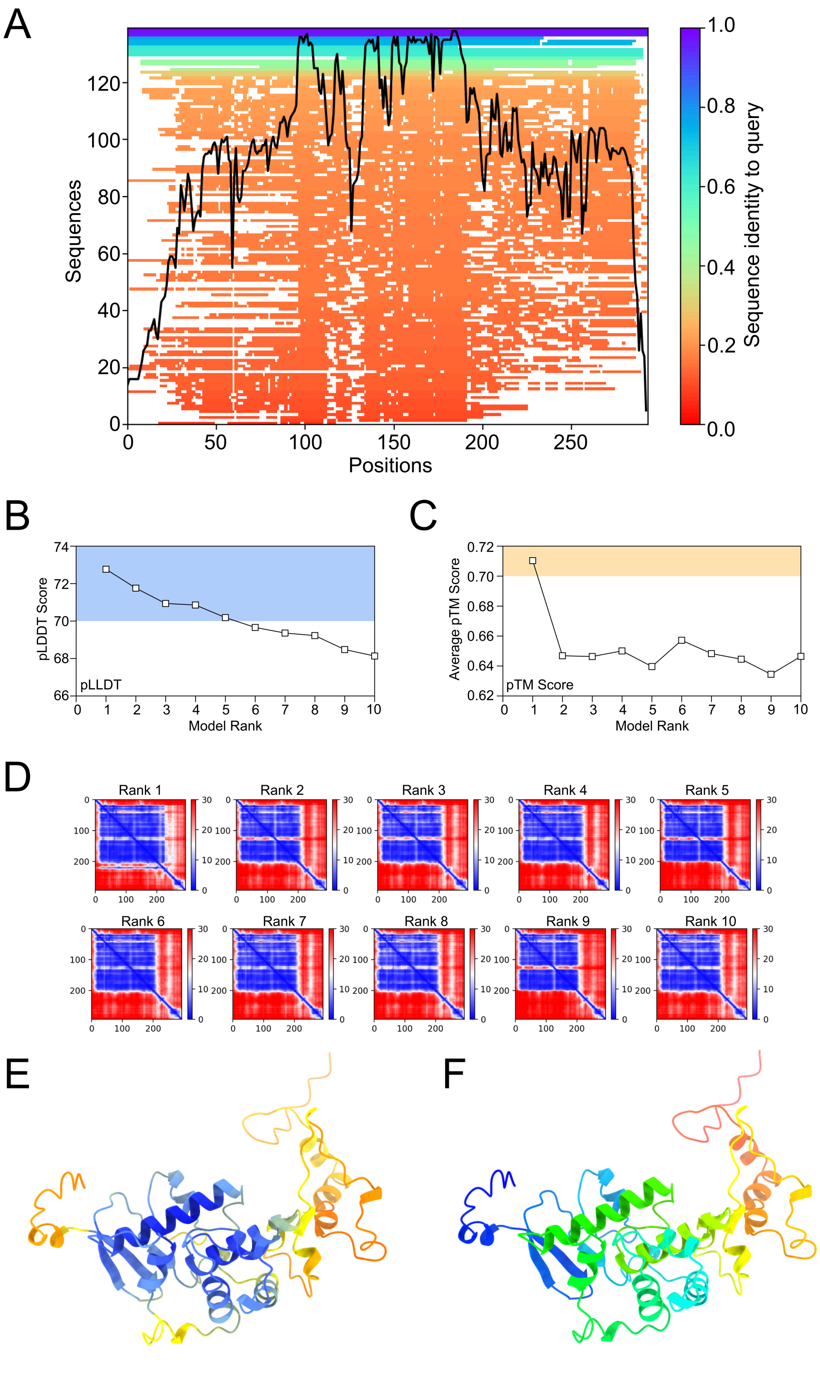


**Fig. S6. AlphaFold structural prediction confidence metrics for SFTSV NSs protein.**

**(A)** Multiple sequence alignment heatmap for SFTSV NSs protein (NCBI Protein: AJD86041) and homologs. Colour scale represents sequence identity from low (red, 0.0) to high (blue, 1.0) conservation. Black line shows position-specific conservation scores. **(B)** Per-residue confidence (pLDDT) scores across AlphaFold model ranks. Blue shading highlights highest confidence region. **(C)** Inter-domain positioning confidence (pTM) scores across model ranks. Yellow shading indicates optimal predictions. **(D)** Predicted Aligned Error (PAE) matrices for top 10 ranked models showing residue-residue distance confidence. Colour scale represents predicted error (0-30 Ångströms), with blue indicating high confidence and red showing uncertainty. **(E)** Rank 1 structure coloured by pLLDT confidence: very high (dark blue: 90-100), confident (light blue: 70-89), low (yellow: 50-69), very low (orange: 0-49). **(F)** Rank 1 structure with rainbow colouring from N-terminus (blue) to C-terminus (red).

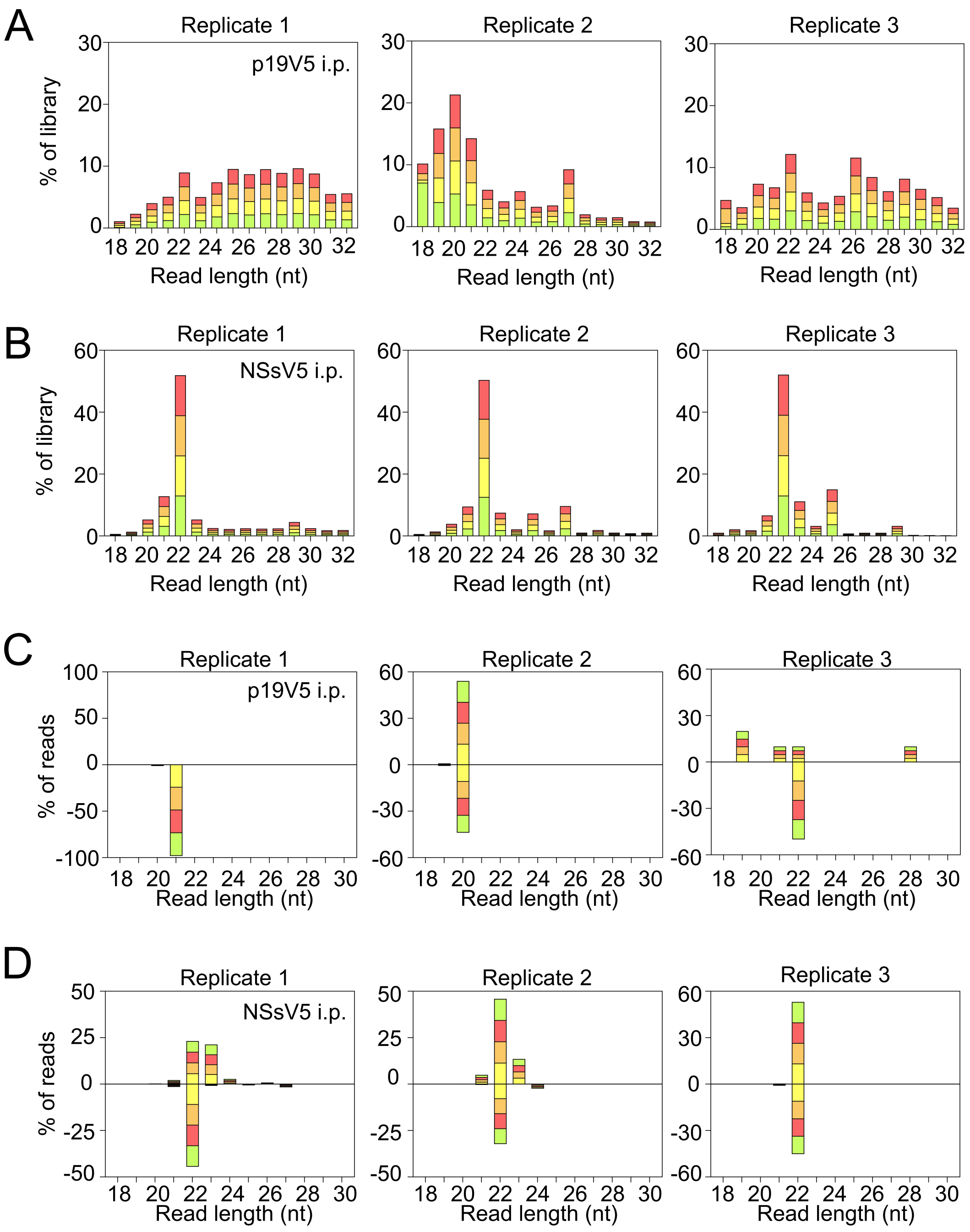


**Fig. S7. Total mapped reads and strand-specific reads for each replicate of Fig. 6B,C.**
Histogram of all small RNA reads (18-30 nt) in samples immunoprecipitated with anti-V5 antibodies at 2 dpi from **(A)** p19V5 or **(B)** NSsV5 SFTSV-infected BME/CTVM6 cells (Replicates 1-3). **(C,D)** Small RNA reads from individual replicates mapped to SFTSV antigenomic (positive values) and genomic (negative values) strand RNAs from **(C)** p19V5 and **(D)** NSsV5 immunoprecipitation samples. Read counts are coloured by 5′ nucleotide identity and grouped by read length. Data are shown as percentage of total mapped reads for each individual biological replicate (n=3).

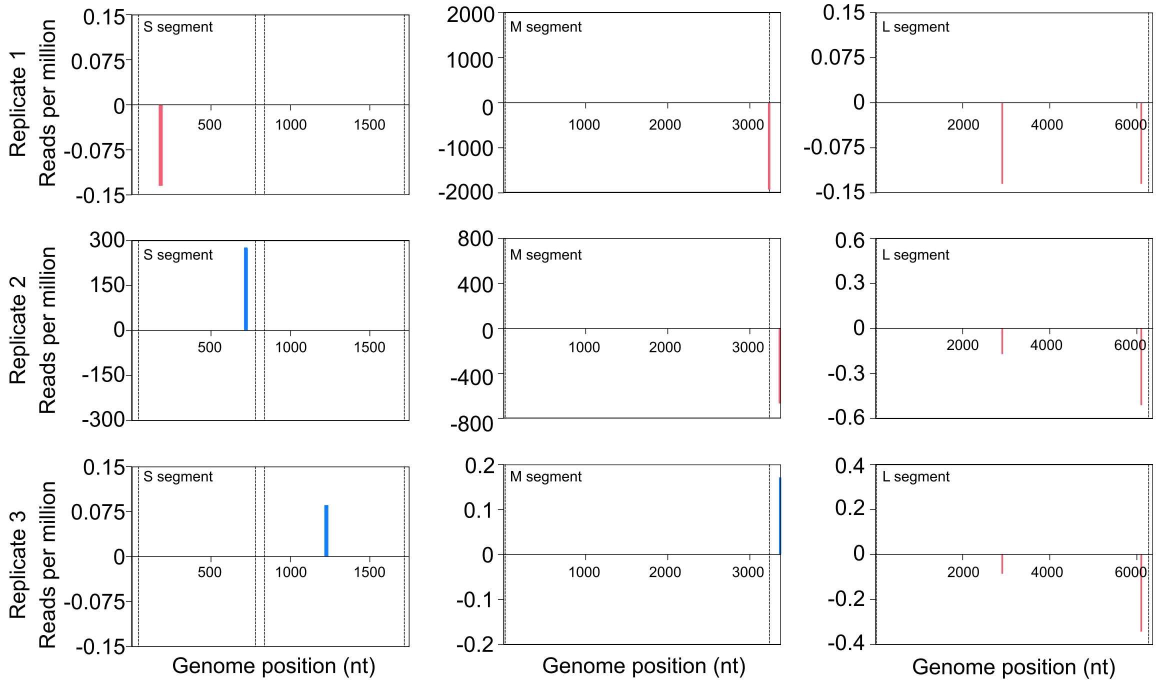


**Fig. S8. Mapping of 22-nt SFTSV-derived vsiRNAs for each replicate of Fig. 6E.**
Mapping of 22-nt SFTSV-derived vsiRNAs across the S (left panel), M (middle panel), and L (right panel) genome segments in p19V5 immunoprecipitation samples (replicates 1-3; reads per million). Data show mapping to the SFTSV antigenomic (blue) or genomic (pink) strands with nucleotide positions indicated. Individual biological replicates (n=3) are shown as mapped read counts at each nucleotide position.

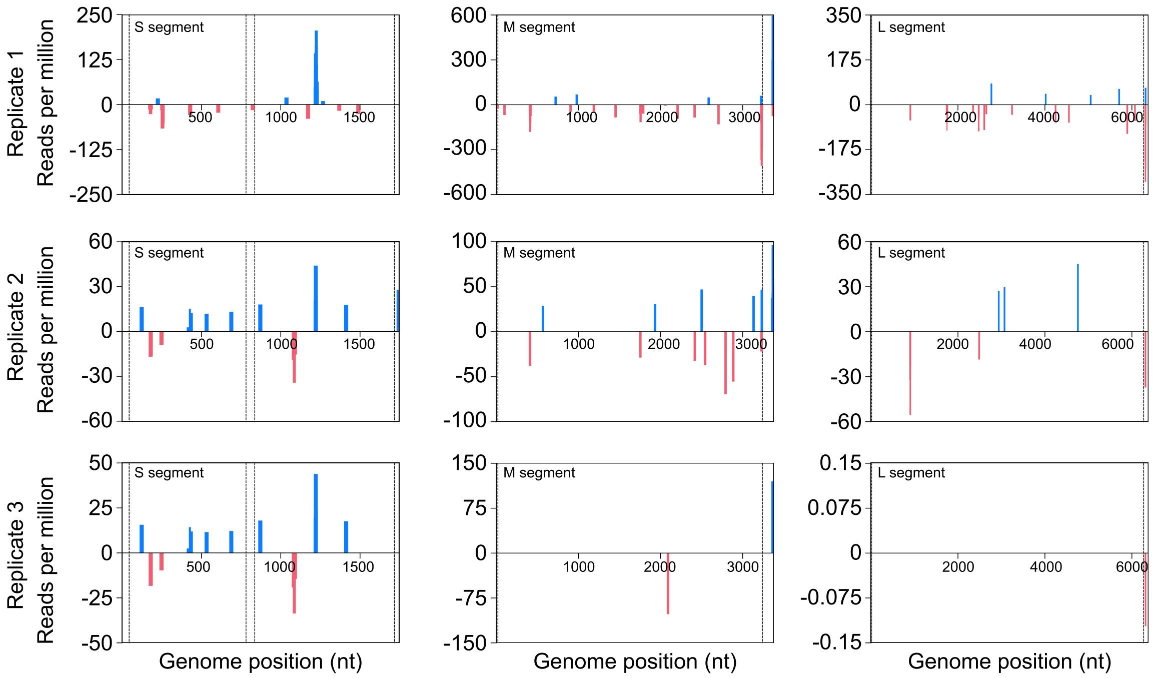


**Fig. S9. Mapping of 22-nt SFTSV-derived vsiRNAs for each replicate of Fig. 6F.**
Mapping of 22-nt SFTSV-derived vsiRNAs across the S (left panel), M (middle panel), and L (right panel) genome segments in NSsV5 immunoprecipitation samples (replicates 1-3; reads per million). Data show mapping to the SFTSV antigenomic (blue) or genomic (pink) strands with nucleotide positions indicated. Mapped read counts at each nucleotide position are shown for individual biological replicates (n=3).

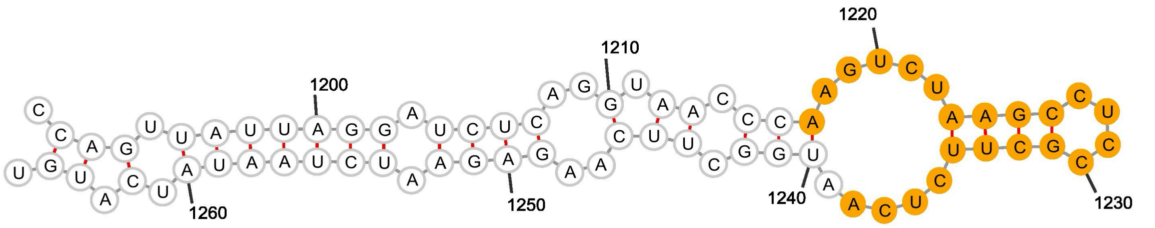


**Fig. S10. Secondary structure of the SFTSV antigenomic S RNA (HB29 strain) showing mapping sites of vsiRNAs immunoprecipitated with NSs from infected cells.**

Secondary structure prediction of a 76-nucleotide sequence (positions 1191-1266) from the antigenomic sense SFTSV S segment RNA (HB29 strain, GenBank: KP202165). Structure was predicted using minimum free energy folding calculations with Turner energy parameters at 37°C via the Forna web server powered by the ViennaRNA package. Orange circles indicate 22-nucleotide siRNAs identified through NSs immunoprecipitation and small RNA sequencing. The distribution of NSs-bound siRNAs across stem-loop and single-stranded regions demonstrates the RNA-binding specificity of NSs and supports its function as a viral suppressor of RNA silencing that sequesters host siRNAs.

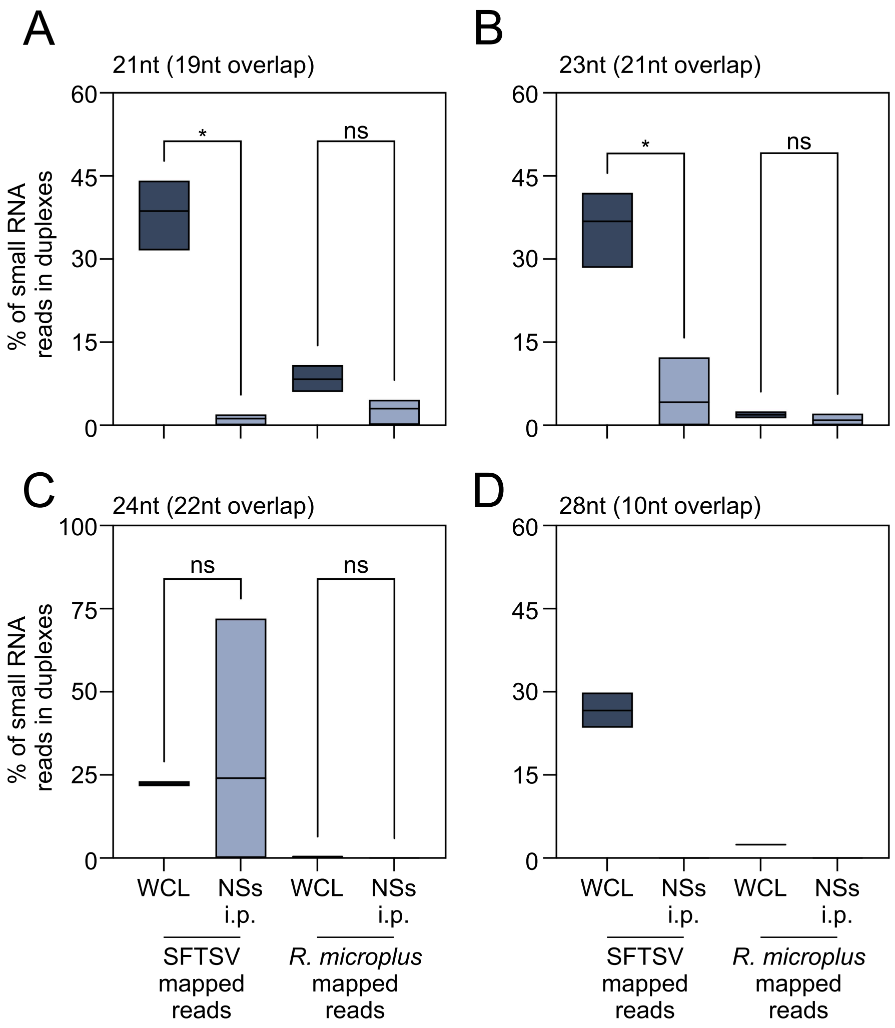


**Fig. S11. Additional detected RNA duplexes of differing sizes in immunoprecipitated samples from Fig. 6.**

Histograms showing the percentage of small RNA reads in duplexes with complementary overlaps: **(A)** 21-nt reads with 19-nt overlap, **(B)** 23-nt reads with 21-nt overlap, **(C)** 24-nt reads with 22-nt overlap, and **(D)** 28-nt reads with 10-nt overlap. Reads were mapped to SFTSV genome or *R. microplus* genome (GCA_002176555.1) in whole cell lysate (WCL) and NSsV5 immunoprecipitation (i.p.) samples. Data represent n=3 biological replicates. Statistical significance was determined using Brown-Forsythe and Welch's ANOVA with Dunnett's T3 multiple comparison test. Asterisks indicate significance: * p ≤ 0.05, ** p ≤ 0.01; ns = not significant.

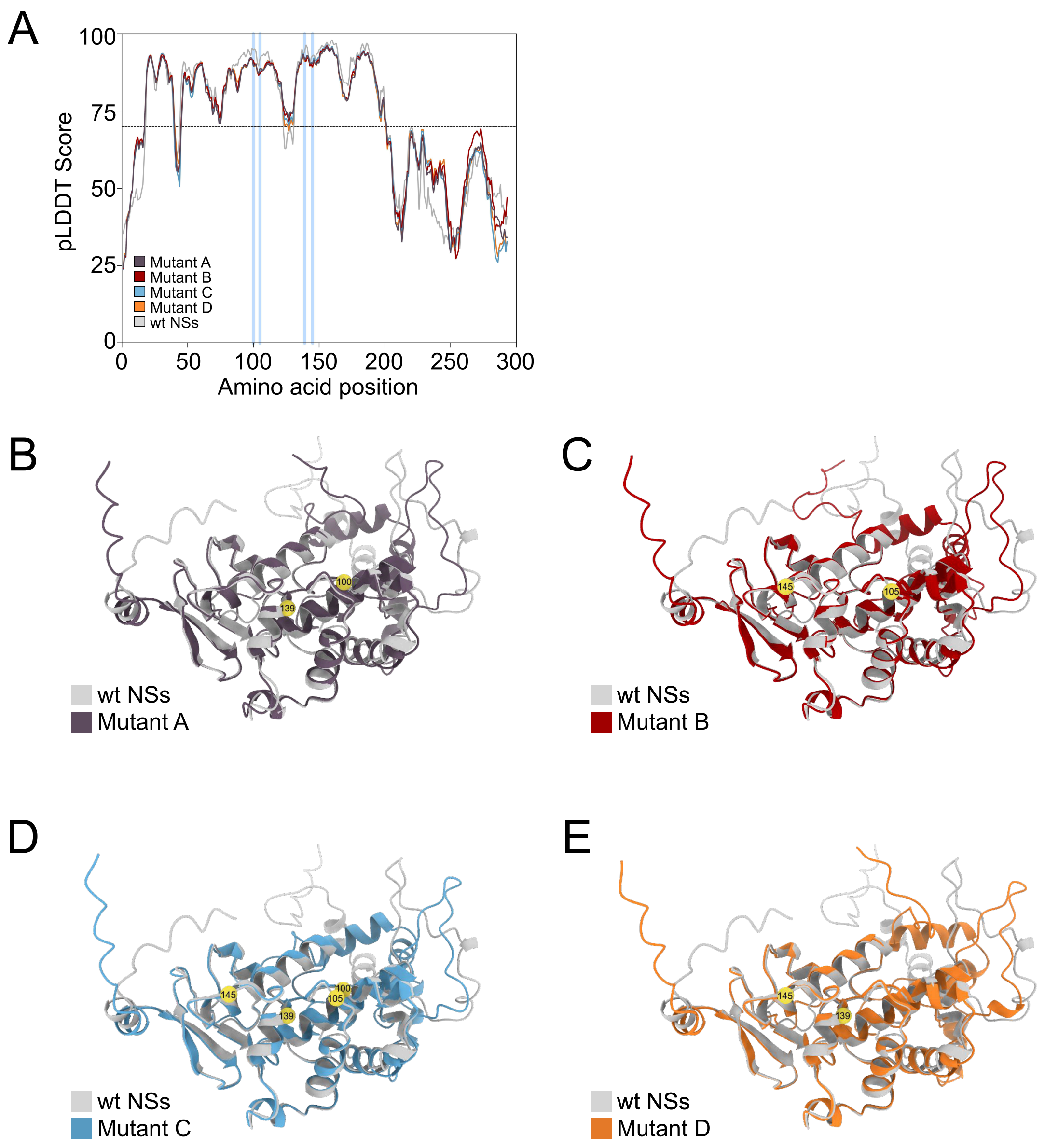


**Fig. S12. Alphafold3-predicted structural models of SFTSV NSs charge-reversal mutants.**
**(A)** Per-residue confidence (pLDDT) scores across AlphaFold models of SFTSV NSs charge mutants. Dashed line indicates high confidence region [>70 score]. Blue bars indicate amino acid positions 100, 105, 139 & 145, that are altered in the NSs mutants. **(B-E)** Predicted structures of NSs showing overlays of wild-type (wt) NSs (light grey) with individual mutants. **(B)** Mutant A, **(C)** Mutant B, **(D)** Mutant C, and **(E)** Mutant D are shown in distinct colours as indicated. Residues targeted for mutation are highlighted and labelled in yellow in each model.

#### Tables

**Table S1. RNA-binding probabilities in SFTSV NSs identified by BindUP.**
Data table reporting the output of the BindUP analysis used to colour Fig. 5D.

| Patch | Amino Acid Residues |
|---|---|
| Patch 1 | LYS51 PHE63 TYR85 ASP87 MET88 ARG89 SER92 GLN93 ARG94 LEU96 SER97 ALA98 ARG100 TRP101 PRO102 SER103 GLY104 LYS105 PRO106 SER107 VAL108 TRP109 GLN112 ALA113 MET116 PHE117 ILE120 ASN122 ALA125 MET126 ARG130 GLU131 ASN132 ARG134 GLY135 LEU136 HIS138 ARG139 ILE140 THR141 LYS142 GLY143 GLN144 LYS145 GLU149 MET152 GLU158 ALA159 GLU161 LYS162 ARG163 LEU165 ARG166 LEU167 GLY168 LEU186 SER190 ARG193 LEU194 ARG195 |
| Patch 2 | MET1 SER2 LEU3 SER4 LYS5 CYS6 SER7 ASN8 |
| Patch 3 | ARG195 PRO222 VAL223 ARG224 LYS225 LYS226 LYS227 THR228 ASP229 TYR232 SER233 VAL234 LEU235 ASP236 LEU244 |

**Table S2. Abundance and sequence identity of vsiRNAs immunoprecipitated with SFTSV NSs.**
* Indicates RNA was used in structural modelling vsiRNA with SFTSV NSs; Target refers to individual SFTSV genome segments. Strands: (-) genomic sense, (+) antigenomic sense.

| Target | Abundance | Mapping counts | Start Position | End Position | Strand | Sequence |
|---|---|---|---|---|---|---|
| wt S (NSsV5) | 1 | 1154 | 245 | 266 | - | ATCGTCAAGGCATCAGGGAAAA |
| | 2 | 1091 | 1211 | 1232 | + | TAACCCAAGTCTAAGCCTCCGC |
| | 3 | 1071 | 1210 | 1231 | + | GTAACCCAAGTCTAAGCCTCCG |
| | 4* | 1070 | 1212 | 1233 | + | AACCCAAGTCTAAGCCTCCGCT |
| | 5 | 927 | 1217 | 1238 | + | AAGTCTAAGCCTCCGCTTCTCA |
| | 6 | 886 | 1162 | 1183 | - | GGAGGGCCACATCCAGAATTGG |
| | 7 | 658 | 419 | 440 | - | CTCCCAGTAGGGCCAGCCGTCA |
| | 8 | 593 | 1479 | 1500 | - | AACTGCTCAAGTCCGGGCTCAA |
| | 9 | 518 | 1216 | 1237 | + | CAAGTCTAAGCCTCCGCTTCTC |
| | 10 | 499 | 596 | 617 | - | ATGCGCGGAGCCAGCAAGACAG |
| M | 1* | 8354 | 3357 | 3378 | + | AAGTGTTGGCCGGTCTTTGTGT |
| | 2 | 8354 | 3357 | 3378 | - | AAGTGTTGGCCGGTCTTTGTGT |
| | 3 | 6811 | 3356 | 3377 | + | GAAGTGTTGGCCGGTCTTTGTG |
| | 4 | 5608 | 3222 | 3243 | - | GAGGACGAAGCTGGCTTAGATG |
| | 5 | 2945 | 2696 | 2717 | - | ATTATAGAGGCCTTCGATTAAG |
| | 6 | 2146 | 3219 | 3240 | - | ATCGAGGACGAAGCTGGCTTAG |
| | 7 | 2089 | 2198 | 2219 | - | CATGGGTCCCATCAGCAGTTAT |
| | 8 | 1909 | 2405 | 2426 | - | CTGGCATGGGGGTTGTAGGCAA |
| | 9 | 1894 | 1443 | 1464 | - | GGCAATTGGGGTCTGGGGATCA |
| | 10 | 1714 | 406 | 427 | - | TTTGAACGGCCAACTTTTGATG |
| L | 1 | 3901 | 6303 | 6324 | - | GTCTGTGGGTGGCTAGGGAGTG |
| | 2 | 3199 | 6305 | 6326 | - | CTGTGGGTGGCTAGGGAGTGTT |
| | 3* | 3199 | 6305 | 6326 | + | CTGTGGGTGGCTAGGGAGTGTT |
| | 4 | 2484 | 5885 | 5906 | - | CTGGAGGAGGACATTGACTTTT |
| | 5 | 2195 | 2596 | 2617 | - | CTACTCAGAAGTCCAGACCAAA |
| | 6 | 1848 | 2760 | 2781 | + | AGCATGGTGGCCTCAGAGAGAT |
| | 7 | 1544 | 4543 | 4564 | - | GCGGTGCAAGGCAGAGGATCTG |
| | 8 | 1534 | 4236 | 4257 | - | GGGTGGTGGCAGCAGGAGTATA |
| | 9 | 1399 | 6061 | 6082 | - | GGACCAAGCTGCCATCACAATG |
| | 10 | 1380 | 5698 | 5719 | + | GTCACATGCCTCAGTTCTCCTG |

**Table S3. Sequencing and library statistics for small RNA beta-elimination study (size range 18-30 nt) used in this study.**

| Sample | Replicate | SRA Accession | Reads in library | After trimming | Mapping to viral genome | % all reads mapping to genome | 22nt Mapping | % of 22nt mapped reads |
|---|---|---|---|---|---|---|---|---|
| Inf3d | 1 | SRR34822371 | 25,270,660 | 25,052,374 | 84,839 | 0.33865 | 84,037 | 0.318098 |
| | 2 | SRR34822370 | 25,272,643 | 24,977,887 | 79,294 | 0.31746 | 79,691 | 0.336446 |
| | 3 | SRR34822359 | 25,272,849 | 24,487,279 | 46,885 | 0.19147 | 46,544 | 0.325438 |
| Inf6d | 1 | SRR34822348 | 25,203,593 | 24,308,520 | 144,414 | 0.59409 | 140,217 | 0.191472 |
| | 2 | SRR34822337 | 25,392,392 | 25,060,705 | 51,523 | 0.20559 | 46,763 | 0.559509 |
| | 3 | SRR34822334 | 25,278,221 | 23,734,717 | 172,166 | 0.72538 | 163,801 | 0.197024 |
| Mock3d | 1 | SRR34822333 | 25,376,299 | 24,489,173 | 240 | 0.00098 | 24 | 0.668871 |
| | 2 | SRR34822332 | 25,253,308 | 24,872,201 | 149 | 0.00060 | 11 | 0.000096 |
| | 3 | SRR34822331 | 25,332,618 | 25,204,144 | 2 | 0.00001 | - | 0.000044 |
| Mock6d | 1 | SRR34822330 | 25,282,979 | 22,387,098 | 23 | 0.00010 | - | 0.000000 |
| | 2 | SRR34822369 | 25,355,105 | 24,208,018 | 19 | 0.00008 | - | 0.000000 |
| | 3 | SRR34822368 | 25,282,133 | 24,501,391 | 12 | 0.33768 | 1 | 0.000000 |
| AF525SFV | 1 | SRR34822354 | 25,309,559 | 25,040,007 | 3,450,730 | 14.08381 | 91,680 | 0.000004 |
| | 2 | SRR34822353 | 25,332,022 | 10,971,923 | 3,080,175 | 12.30101 | 80,731 | 0.835587 |
| | 3 | SRR34822352 | 25,319,300 | 25,090,446 | 2,708,472 | 24.68548 | 80,516 | 0.321760 |
| AF525Mock | 1 | SRR34822351 | 25,346,519 | 25,047,968 | 990 | 0.00395 | 17 | 0.321447 |
| | 2 | SRR34822350 | 25,295,872 | 25,197,970 | 64 | 0.00026 | - | 0.000067 |
| | 3 | SRR34822349 | 25,303,505 | 25,098,903 | 35 | 0.00014 | 2 | 0.000000 |
| Inf3dB | 1 | SRR34822367 | 25,350,178 | 24,907,717 | 726 | 0.00289 | 150 | 0.000008 |
| | 2 | SRR34822366 | 25,252,985 | 24,635,793 | 485 | 0.00195 | 71 | 0.000609 |
| | 3 | SRR34822365 | 25,331,899 | 25,012,897 | 230 | 0.00093 | 43 | 0.000284 |
| Inf6dB | 1 | SRR34822364 | 25,425,120 | 25,189,475 | 1,786 | 0.00714 | 337 | 0.000171 |
| | 2 | SRR34822363 | 25,376,081 | 25,188,891 | 4,400 | 0.01747 | 3,993 | 0.001338 |
| | 3 | SRR34822362 | 25,247,125 | 25,211,477 | 3,642 | 0.01446 | 2,296 | 0.015838 |
| Mock 3dB | 1 | SRR34822361 | 25,356,363 | 25,080,844 | 429 | 0.00170 | 70 | 0.009154 |
| | 2 | SRR34822360 | 25,358,985 | 25,205,390 | 20 | 0.00008 | 4 | 0.000278 |
| | 3 | SRR34822358 | 25,277,402 | 25,186,702 | 22 | 0.00009 | - | 0.000016 |
| Mock 6dB | 1 | SRR34822357 | 25,226,225 | 24,800,147 | 25 | 0.00010 | - | 0.000000 |
| | 2 | SRR34822356 | 25,406,194 | 25,105,036 | 63 | 0.00025 | - | 0.000000 |
| | 3 | SRR34822355 | 25,404,511 | 25,240,186 | 63 | 0.00025 | 1 | 0.000000 |
| AF525SFVB | 1 | SRR34822347 | 25,390,903 | 25,175,217 | 340,327 | 1.34835 | 1,389 | 0.000004 |
| | 2 | SRR34822346 | 25,321,281 | 25,065,359 | 327,499 | 1.30088 | 1,337 | 0.005542 |
| | 3 | SRR34822345 | 25,283,213 | 25,096,582 | 307,466 | 1.22666 | 981 | 0.005327 |
| AF525MockB | 1 | SRR34822344 | 25,249,702 | 25,061,900 | 271 | 0.00108 | 3 | 0.003914 |
| | 2 | SRR34822343 | 25,227,225 | 25,001,085 | 248 | 0.00006 | - | 0.000012 |
| | 3 | SRR34822342 | 25,248,873 | 25,124,411 | 14 | 0.00005 | - | 0.000000 |

**Table S4. Sequencing and library statistics for small RNA immunoprecipitation study (size range 18-30 nt) used in this study.**

| Sample | Replicate | SRA Accession | Reads in library | After trimming | Mapping to viral genome | % all reads mapping to genome | 22nt Mapping | % of 22nt mapped reads |
|---|---|---|---|---|---|---|---|---|
| NSs_2dpi | 1 | SRR34822341 | 25,246,736 | 22,730,371 | 142,519 | 0.62700 | 107,931 | 0.47483 |
| | 2 | SRR34822340 | 25,169,923 | 23,736,457 | 38,291 | 0.16132 | 31,209 | 0.13148 |
| | 3 | SRR34822339 | 25,192,518 | 24,613,751 | 11,997 | 0.04874 | 5,460 | 0.02218 |
| p19_2dpi | 1 | SRR34822338 | 13,938,209 | 7,418,505 | 14,285 | 0.19256 | 3 | 0.00004 |
| | 2 | SRR34822336 | 17,957,002 | 11,739,367 | 11,099 | 0.09455 | 13 | 0.00011 |
| | 3 | SRR34822335 | 18,085,230 | 11,661,667 | 9.0 | 0.00008 | 6 | 0.00005 |

#### Datasets

**Dataset S1. RBDetect output**
Data table reporting the output of the RBDetect analysis used to colour Fig. 5D.

**Dataset S2. RNA-NSs protein interaction data.**
Identification of RNA-binding interfaces and contact residues within 4 Å of vsiRNA substrates, and structural determinants of vsiRNA recognition for 11 small RNAs identified in experimental data from Table S2.
